# Spatial constraints drive amylosome-mediated resistant starch degradation by *Ruminococcus bromii* in the human colon

**DOI:** 10.1101/2025.03.31.646359

**Authors:** Benedikt H Wimmer, Sarah Moraïs, Itai Amit, Omar Tovar-Herrera, Meltem Tatli, Anke Trautwein-Schult, Paloma Tödtli, Sebastian Simoni, Matteo Lisibach, Liron Levin, Dörte Becher, Edward A. Bayer, Ohad Medalia, Itzhak Mizrahi

## Abstract

Degradation of complex dietary fiber by gut microbes is essential for colonic fermentation, short-chain fatty acid production, and microbiome function. *Ruminococcus bromii* is the primary resistant starch (RS) degrader in humans, which relies on the amylosome, a specialized cell-bound enzymatic complex. To unravel its architecture, function, and the interplay among its components, we applied an holistic multilayered approach and found that amylosome composition RS degradation, and enzymatic synergy are regulated at two levels: structural constraints enforcing enzyme proximity and expression-driven shifts in enzyme proportions. Cryo-electron tomography revealed that the amylosome comprises a constitutive extracellular layer extending toward the RS. However, proteomics demonstrated its remodeling across different growth conditions, with Amy4 and Amy16 comprising 60% of the amylosome in response to RS. Structural and biochemical analyses revealed complementarity and synergistic RS degradation by these enzymes, which allow *R. bromii* to fine-tune its adaptation to dietary fiber and shape colonic metabolism.

## Introduction

Dietary fiber is essential for promoting a diverse and balanced human microbiome, which in turn enhances metabolic function, reduces inflammation, and protects against chronic diseases^1,2^. Dietary fibers are composed of various carbohydrates that are indigestible to the human host and remain structurally intact until they reach the lower gastrointestinal tract, where they undergo microbial fermentation. Dietary fiber carbohydrates are degraded by specialized microbes into simple sugars fueling the entire microbial community^3^. This process plays a crucial role in microbiome-host interactions, by producing metabolites such as short-chain fatty acids (SCFAs), which promote host health^4^.

Resistant starch (RS) is a major dietary fiber component, whose fermentation has been linked to various health benefits, including improved insulin sensitivity^5^, reduced risk of colorectal cancer^6^, protection against infectious diarrhea^7^, reduced inflammation in chronic kidney disease^8^, and enhanced glucose control and satiety^9^. RS typically forms large, insoluble granules that, like other dietary fibers, cannot be degraded by human enzymes. Instead, its breakdown relies entirely on microbial fermentation, during which RS is converted into simpler sugars that fuel both RS-degrading bacteria and secondary consumers in the gut microbiome.

To date, only a few bacterial species have been identified as RS degraders, the major species being *Ruminococcus bromii* and *Bifidobacterium adolescentis*^10^. Among them, *R. bromii* is the dominant degrader, accounting for 80% of RS-adherent bacteria in human fecal samples^11^ and outcompeting *B. adolescentis* on a wide range of recalcitrant substrates^12^. This specialization positions *R. bromii* as the keystone species for RS degradation in the gut^12^, enabling metabolic cross-feeding that supports the growth of other microbiota members, such as *Bacteroides thetaiotaomicron*^13^ and *Ruminococcus gnavus*^14^. *R. bromii* abundance is associated with multiple health benefits, including enhanced response to cancer immunotherapy^15^, protection against chronic kidney disease^16^, and reduced risk of depressive disorders^17^. Some of these benefits stem from its ability to stimulate production of SCFAs, particularly butyrate^18^, a key regulator of immune function and inflammation^4^.

The ability of *R. bromii* to efficiently deconstruct RS while simultaneously supporting the broader gut ecosystem is attributed to a cell-adjacent protein complex known as the amylosome^19^. While early studies inferred the potential role of this complex in RS degradation using *R. bromii* cell extracts, the precise function of its components and how they interact with RS remain unclear. Structurally, the amylosome resembles the well-characterized cellulosome^20^, exhibiting a high degree of adaptability. It consists of two protein groups: (i) the scaffoldins, which contain cohesin domains and serve as structural organizers, either attaching to the cell wall or remaining free in the extracellular space, and (ii) the dockerin-containing proteins, which bind to the scaffoldins via specific cohesin-dockerin interactions and include key carbohydrate-active enzymes (CAZymes)^21^.

Genomic analyses identified five scaffoldins (Sca1–Sca5) and 27 dockerin-containing proteins in *R. bromii*^19,22^. Among them, five dockerin-containing proteins—Amy4, Amy9, Amy10, Amy12, and Amy16—also contain CAZyme domains, rendering them primary candidates for RS degradation. One enzyme, Amy4 (also known as Sca1), exhibits a unique dual-domain architecture, possessing both a cohesin and a dockerin domain, thereby functioning as both a scaffoldin and a GH13-family α-amylase. This structural duality suggests that Amy4 plays a central role in amylosome assembly and function^19^. The ability of Amy4 to homo-polymerize through intermolecular cohesin-dockerin interactions may further contribute to amylosome organization and enzyme recruitment. The scaffoldins Sca2 and Sca5 have a sortase recognition motif, indicating that they are anchored to the cell wall, while Sca3 and Sca4 are presumed to be in the cell-free state. Among the dockerin-containing proteins, the five amylosomal enzymes underwent preliminary characterization for their substrate preferences: Amy4, Amy9, and Amy16 exhibited amylase activity, while Amy10 and Amy12 were specific for pullulan degradation^22^. Amy10 and Amy12 were further characterized, indicating that they accumulate maltotriose from pullulan, and revealing the mechanism of action for Amy12^23^. Key carbohydrate-binding domains of the starch-adherence systems, Sas6^24^ and Sas20^25^, were also characterized for their structure and binding behavior. Sas20 is especially notable, as it is consistently among the most highly expressed amylosome proteins and binds exclusively and uniquely to cohesin domain 6 of Sca5^25^.

Despite these insights, the structural organization, functional interactions between the enzymes, and precise role of the amylosome in RS degradation remain largely unexplored. In this study, we therefore aimed to elucidate the molecular architecture of the amylosome, identify key enzymes involved in RS degradation, connect their structure to function, determine how its components interact, and uncover the mechanistic basis for *R. bromii’*s exceptional ability to degrade RS.

We leveraged *in situ* cryo-electron tomography (cryo-ET) to visualize the architecture of the amylosome, revealing it as a densely packed extracellular layer extending from *R. bromii* toward the RS substrate. This organization reinforces the concept of a highly structured, cell-bound degradation system. Next, quantitative proteomics demonstrated that amylosome composition shifts dynamically in response to different growth stages and carbon sources, allowing us to identify its major CAZymes. To further elucidate their function, we applied cryo-EM to resolve the structure of four key amylosome enzymes and identify structural variations in their active sites, providing insights into their substrate preferences and potential functional complementarity. Biochemical assays and interaction studies identified Amy4 and Amy16 as key RS-degrading enzymes and revealed that they exhibit strong synergistic activity in RS breakdown, a synergy critically dependent on their spatial organization. Notably, while the amylosome exhibits high combinatorial flexibility, we found that the degrees of freedom are constrained in a manner that directs Amy4 and Amy16 into close proximity, ensuring efficient RS degradation. Taken together, our findings provide a comprehensive physiological, structural, and functional understanding of amylosome-mediated RS metabolism by *R. bromii*, a keystone species of the gut microbiome.

## Results

### Amylosomes appear as dense, constitutively expressed structures at the periphery of *R. bromii* cells

To elucidate *R. bromii*’s ability to degrade RS via its amylosome system, we first characterized the spatial organization of amylosomes around *R. bromii* cells, using fluorescence imaging, and cryo-ET. We visualized amylosome distribution around bacteria grown on either fructose, the simplest carbon source that *R. bromii* can utilize, or RS, where *R. bromii*’s role as a keystone species in the human gut microbiome is manifested. Our results reveal that amylosomes form dense structures anchored to the cell wall on almost all cells, even in the absence of RS (Fig. 1, Fig. S1a).

**Figure 1:**
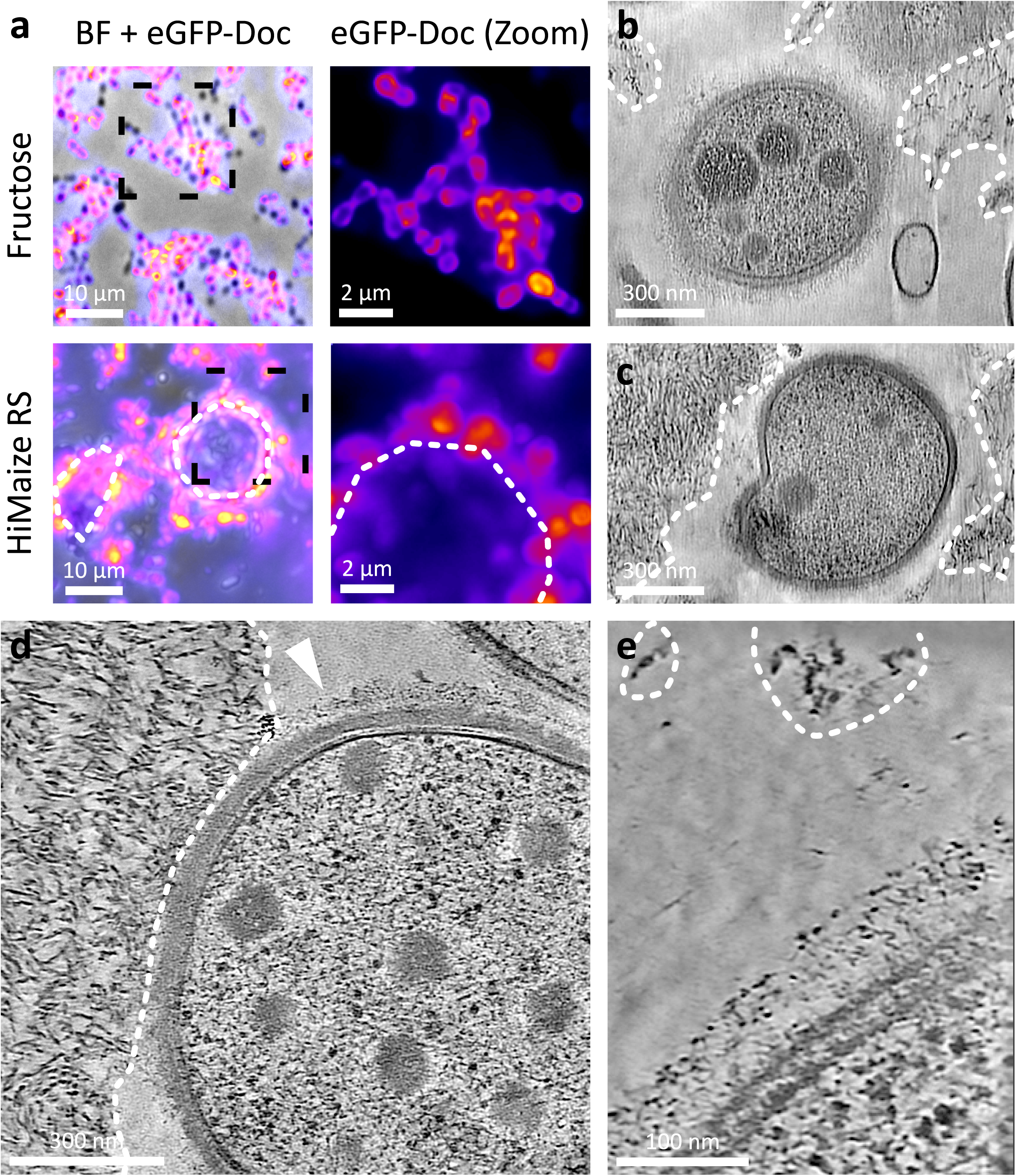
Amylosomes appear as dense, constitutively expressed structures at the periphery of *R. bromii* cells. **(a)** Staining *R. bromii* with an eGFP-Doc13a, an amylosome label shown here in magenta, reveals that almost all cells seen in brightfield (BF) imaging (grey) express amylosomes, both after growth on fructose and HiMaize resistant starch. A zoom-in (right) clearly shows that the staining is localized to the cell surface. Here, only the fluorescence signal is shown. **(b)**, **(c)** Crops from a low-magnification tomogram of *R. bromii* grown on RS granules. Bacterial cells are adhering to the RS substrate with a maximal distance of 200 nm. **(d)** At higher magnifications, direct contacts between bacterium and RS can be seen. The cell is surrounded by a dense protein layer, the putative amylosome (arrow). **(e)** High-magnification tomogram of the edge of *R. bromii* grown on RS shows the cells in close proximity with the substrate. The architecture of the amylosomes is visible as globular densities anchored to the cell wall through elongated linkers. Resistant starch granules are outlined with a dashed line in all panels.

To assess the prevalence of amylosomes on a population level, we used light and fluorescence microscopy and labelled cells with a previously described GFP-dockerin probe that binds to exposed cohesin domains on the cell surface^22^. Fluorescence imaging confirmed that amylosomes are detected around almost all *R. bromii* cells, forming a distinct rim around the cell, regardless of whether fructose or RS was supplied as the carbon source (Fig. 1a). To further resolve the molecular architecture of amylosomes, we applied cryo-focused ion beam (FIB) milling followed by cryo-ET to vitrified *R. bromii* cultures. Low-magnification cryo-ET revealed that bacterial cells are embedded in the outer layer of RS granules, with distances of up to 200 nm between the cell wall (CW) and the nearest RS surface (Fig. 1b,c). Higher magnifications showed direct cell-substrate contact at specific sites, with an extracellular protein layer extending from the CW (Fig. 1d). These extracellular densities were observed in 85% (111/130) of fructose-grown cells and 90% (133/148) of RS-grown cells, closely matching the proportion of amylosome-positive cells identified through fluorescent labelling. The layer structurally resembled the *Clostridium thermocellum* cellulosome^20^, supporting its identification as the amylosome.

Cryo-ET analysis enabled nanometer-scale 3D analysis of *R. bromii’*s extracellular architecture. High-magnification tomograms revealed amylosomes as a dense protein layer attached to the CW through string-like protrusions, both in RS-grown cells (Fig. 1e) and fructose-grown cells (Fig. S1a), extending toward the substrate after growth on RS. We focused our analysis on a subset of 21 tomograms (10 RS- and 11 fructose-grown cells), in which both cytoplasmic and extracellular structures were clearly resolved and the cell membrane (CM) and CW were contained in cross-section. To analyze the *in situ* architecture of the amylosome, we used a blob-picking approach to naively select all extracellular protein densities. This revealed no significant differences in amylosome architecture between carbon sources: amylosomes were densest 30–50 nm outside the CW, with 65% of proteins spaced only 8–12 nm apart from their nearest neighbor (Fig. S1b).

To validate molecular preservation, we performed sub-tomogram averaging (STA) of ribosomes in the cytoplasm, achieving a resolution of 14 Å from 1,135 particles (Fig. S2). However, STA and classification of amylosome components could not recover the identities of individual proteins, likely due to the heterogeneity and relatively low molecular weights of amylosome-associated proteins (37 proteins, molecular weights: 10 - 150 kDa).

As we observed no significant changes in the prevalence or architecture of the amylosome between cells grown on fructose and RS, we hypothesized that the enzymatic composition of the amylosome would be dynamically regulated, even though the overall structure remains unchanged. This could allow dynamic fine-tuning degradative capabilities in response to changing environmental cues.

### Carbon source-driven remodeling of amylosome composition

To test our hypothesis, that *R. bromii* adapts to changing carbon sources by modulating amylosome composition, and to identify key proteins that vary in their proportions and are important for *R. bromii* amylosome function, we conducted a proteomics analysis of bacteria grown on four carbon sources: RS, fructose, soluble starch, and pullulan, which mimics RS branching points. We incorporated a temporal dimension to the proteomic analyses by sampling cultures at three growth phases—mid-logarithmic, late-logarithmic, and stationary (Fig. S3). Given that amylosomes are structurally well-organized around the *R. bromii* cell wall and previous studies have identified amylosome components in both cell-associated and supernatant fractions^19^, we separately analyzed the cell-associated and secreted proteomes.

The overall number of detected proteins varied significantly with the carbon source but was largely unaffected by growth phase (Fig. 2a; two-way ANOVA, 90% vs. 2% variance explained). Principal coordinates analysis (PCoA) of the presence-absence proteome (Jaccard matrix) revealed that the carbon source has a marked influence on both the cell-associated and supernatant proteome (Fig. S4a). RS- and pullulan-grown cultures expressed proteomes divergent from those on fructose or starch-grown cultures. While a broad set of proteins was expressed in cultures grown on simple sugars like fructose and similarly on soluble starch, significantly smaller protein subsets were expressed in RS- and pullulan-grown cultures (Fig. 2a, p < 0.0001, Dunnett’s multiple comparisons test). Heatmaps of all amylosome proteins and free CAZymes involved in starch degradation, expressed in both the cell-associated proteome and the secretome, further illustrate the amylosome component versatility across carbon sources (Figure S5).

**Figure 2:**
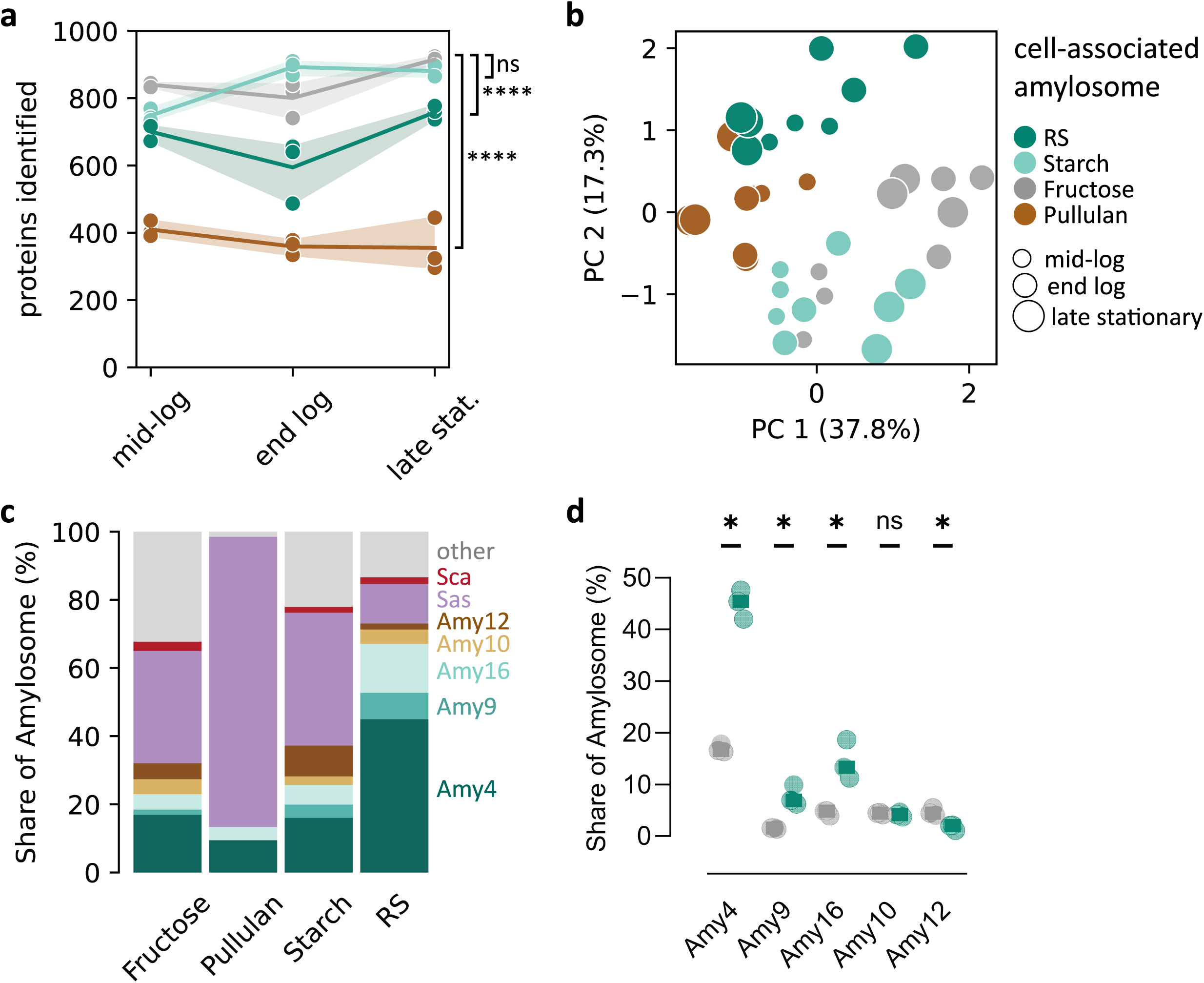
Carbon source-driven remodeling of amylosome composition. **(a)** Depending on the carbon source, significantly different numbers of proteins could be identified, shown here for both fractions together (two-way ANOVA, p < 0.0001). Results from Dunnett’s multiple comparison test between each carbon source and fructose are shown. **(b)** A PCA of the amylosome proteins in the cell-associated proteome shows carbon source-specific adaptation of the amylosome. **(c)** The relative abundance of the different amylosome proteins in the cell-associated amylosome in the late stationary growth phase, across carbon sources. The average abundance from three biological replicates is shown, proteins are colored by their annotated function. **(d)** The relative abundances of the amylosome CAZymes in the cell-associated amylosome after growth on fructose (grey) or RS (dark). Three biological replicates and the results of an unpaired t-test with Holm-Sidak multiple testing correction shown (3 biological replicates).

Our analyses reveal a common core of 329 proteins (proteins identified in 2/3 replicates for each condition). This common core proteome represents 98% of the pullulan expressed proteins, with no proteins specific to growth on pullulan. This suggests that *R. bromii* employs a specialization strategy to adapt protein expression to various complex carbon sources. Among the identified core proteins on RS, 96% were also expressed on starch and fructose, while a set of 20 proteins were specific to growth on RS. Among these 20 RS-specific proteins, 12 were identified in the cell-associated fractions. These included an annotated fructose-1,6-bisphosphatase, part of the gluconeogenesis pathway, and two annotated ATPases, potentially linking RS degradation to human health promotion through cross-feeding and the release of beneficial metabolites.

In our further analysis, we focused on the cell-associated proteome, as the cell-associated fraction harbors substantially higher starch-degrading activity than the supernatant^19^. Principal component analysis (PCA) of the cell-associated protein quantities revealed distinct adaptation patterns of the amylosome proteins (Fig. 2b, ANOSIM by carbon source R = 0.62, p = 0.0001). The abundances of Amy12, Sca2 and Doc17 (a putative protease) are among the strongest contributors to the variance on principal component (PC) 1, while the abundances of Doc8, Doc9 and Doc22 (two uncharacterized proteins and one putative CBM26, respectively), are the strongest contributors to PC2. Furthermore, while Amy4 is consistently the most expressed amylosome enzyme on the cell surface, it is ranked 12th in terms of overall protein expression in RS-grown cells, while in bacteria grown on fructose and starch, Amy4 is ranked 18th and 61st, respectively. The expression of Amy16 is also triggered in RS, where it ranks as the 16th most expressed protein on the cell surface on RS, but only 71st and 132nd most abundant on fructose and starch, respectively. On pullulan, Amy4 and Amy16 rank 33 and 175 most abundant proteins respectively. This suggests a significant modulation of abundance for these two components in RS. Together, these findings further support our hypothesis that amylosome composition is dynamically regulated in response to the substrate. We then focused on identifying amylosome-associated proteins that are fine-tuned based on carbon source availability.

We thus analyzed the share that each amylosome protein occupied at late stationary growth phase for each carbon source (Fig. 2c). This analysis reveals a strong shift in occupancy for the cell-associated amylosome, whereas the cell-free amylosome was remarkably stable in composition (Fig. S4b-d). After growth on fructose, the amylosome occupancy of the cell-associated amylosome is approximately split into thirds: one third consists of the five amylosome CAZymes Amy4, Amy9, Amy10, Amy12, and Amy16, one third of the starch-adherence systems Sas6 and Sas20, and the remainder consists mostly of dockerin-containing proteins with unknown functions. On pullulan, the cell-associated amylosome consists almost exclusively of Sas20, with about 20% Amy4 and Amy16 in the late stationary phase. The pullulanase Amy12 was only detected up to the end-log time point (Fig. S6). On soluble starch, the overall composition is similar to that on fructose, in line with the results of the PCA (Fig. 2b); however, there is a slight increase in the share of CAZymes at the expense of the dockerin-containing proteins of unknown function. On RS, there is a marked departure from the amylosome compositions on all other carbon sources: Amy4 alone accounts for over 40% of amylosome proteins, with Amy16 accounting for another 15% (Fig. 2c). Overall, the CAZymes account for 70% of amylosome proteins. The share of starch-adherence system proteins is drastically reduced. The strong enrichment of CAZyme activity in the amylosome likely reflects the challenge of efficiently degrading the recalcitrant RS substrate. Notably, the CW-anchored scaffoldins Sca2 and Sca5 were not consistently detected in the cell-associated proteome, suggesting they were not efficiently removed during proteomics sample preparation (Fig. S6). However, specific antibodies confirmed their presence on the cell wall using fluorescent immuno-labelling (Fig. S7).

We further wanted to analyze whether the adaptations of the amylosome in cells grown on RS were statistically significant. We thus compared the proportion of each amylosome-associated CAZyme between RS- and fructose-grown cultures. Amy4, Amy9, and Amy16 were significantly enriched from RS-grown cells: Amy4 increased threefold to 45 ± 3%, Amy9 increased fivefold to 8 ± 2%, and Amy16 increased threefold to 14 ± 4%. Meanwhile, Amy10 remained stable at 4 ± 0.4%, whereas Amy12 was depleted 2.5-fold to 1.8 ± 0.5% (Fig. 2d). These insights, revealing that the five enzymatic components of the amylosome are consistently present across all carbon conditions yet dynamically modulated in response to RS, provided a unique opportunity to further investigate both the structure-function relationship of the amylosome and the specific roles of these components in RS deconstruction. The two most abundant amylases Amy4 and Amy16 alone account for 60% of amylosome proteins after growth on RS. We aimed to integrate structure, activity profiles, interactions, and potential enzyme combinatorics to achieve a comprehensive understanding of amylosome function as a whole and its role in RS breakdown in particular.

### Functional and structural characterization of amylosome CAZymes suggests complementary activity in RS decomposition

Building on the constant expression of amylosome components alongside the discrete modulation of specific CAZymes in response to RS, we next sought to investigate *R. bromii*’s hallmark ability to degrade RS. To this end, we assessed the activity against model substrates with glycosidic bonds similar to those found in RS, and we determined the molecular structures of amylosome enzymes Amy4, Amy10, Amy12 and Amy16. Our results revealed distinct substrate specificities and varying promiscuity among the amylosome CAZymes. Notably, Amy4 and Amy16, both predicted to degrade RS, exhibited partially divergent functional properties. Structural analysis of Amy16 revealed a larger active site cavity, consistent with its increased substrate promiscuity. In contrast, Amy4 had a narrower active site, indicative of a more defined substrate specificity.

Due to the structural complexity of RS, we could not directly use it for 3D structure-function analysis. Instead, we tested a range of model substrates with similar glycosidic linkages. RS consists of a complex mixture of amylose — glucose polymers linked via α(1,4)-glycosidic bonds forming helices — with branching points attached through α(1,6)-bonds. This chemical and structural diversity necessitates multiple enzymes for efficient degradation. The five amylosome-associated CAZymes in *R. bromii* share a conserved domain composition: an N-terminal signal peptide for secretion, followed by either a GH13-family amylase (AmyA; α(1,4)-hydrolyzing) or a pullulanase (PulA; α(1,6)-hydrolyzing) domain, along with CBM26 and dockerin domains. The PulA domains are present together with mucin-binding protein (MucBP) domains. The CBM mediates substrate binding^26^, while the dockerin domain enables enzyme-scaffoldin interactions, facilitating amylosome assembly. Additionally, Amy4 carries a cohesin domain, allowing it to recruit other dockerin-containing proteins into the amylosome (Fig. 3a).

**Figure 3:**
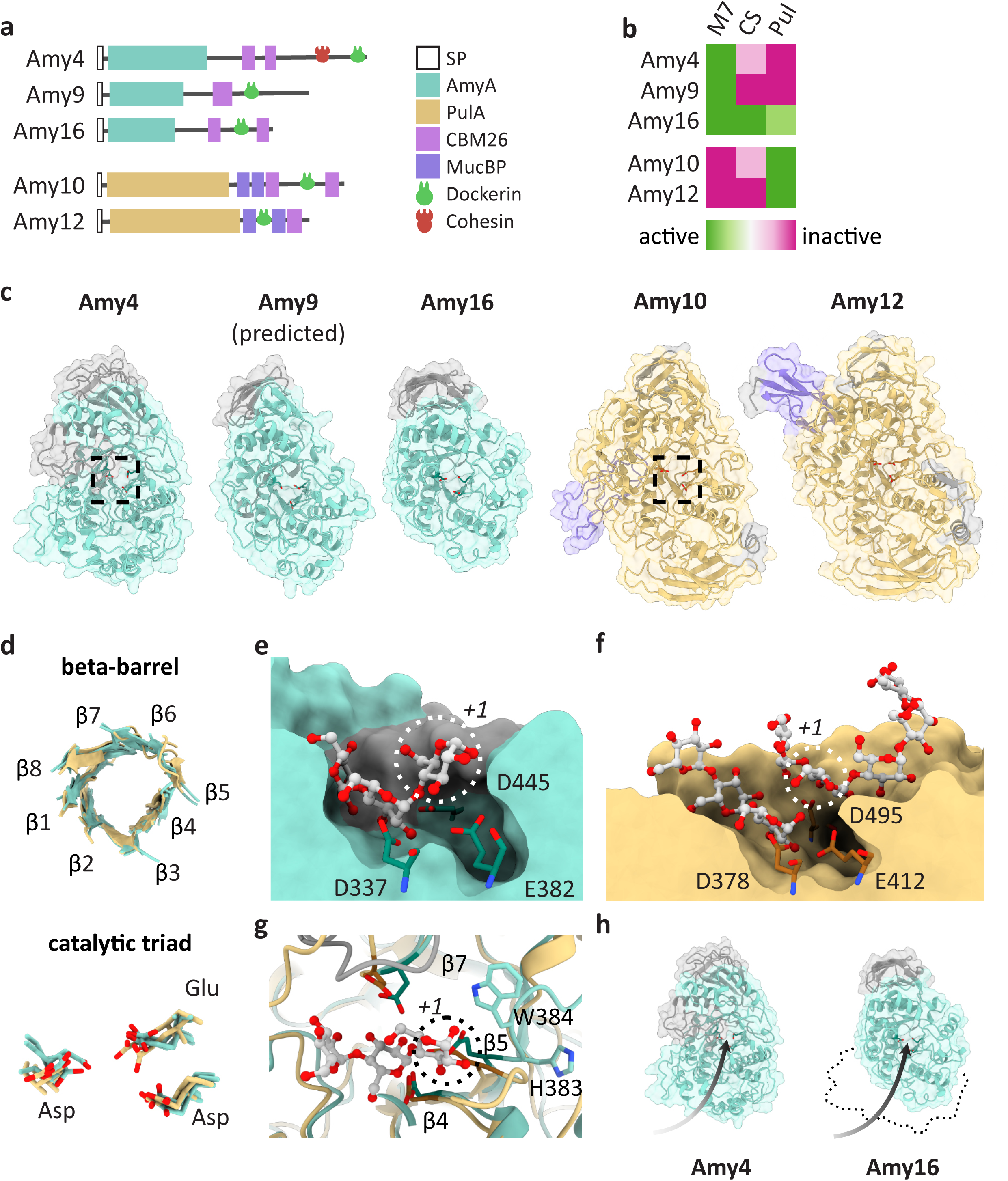
Functional and structural characterization of amylosome CAZymes suggests complementary activity in RS decomposition. **(a)** The five amylosome CAZymes follow a conserved domain order: a signal peptide (SP), followed by the catalytic amylases (AmyA) or pullulanase (PulA) domain, complemented with carbohydrate-binding modules (CBM) or mucin-binding protein domains (MucBP). All contain a dockerin domain, which allows anchoring to the amylosome. Amy4 has an additional cohesin domain, to anchor other enzymes. **(b)** In *in vitro* assays, Amy4 and Amy16 act on (1,4) bonds in maltoheptaose (M7), Amy10 and Amy12 act on (1,6) bonds in pullulan (Pul). Additionally, Amy16 is also active on cornstarch (CS), and shows residual activity against Pul. Activity of Amy4 against CS and of Amy10 against M7, which were previously reported^22,23^, could not be replicated here, but are indicated by a pink shade. **(c)** The overall molecular structures of the amylosome CAZymes: Amy4, Amy9, Amy16, Amy10 and Amy12. The structures for Amy4, Amy10, Amy12 and Amy16 were experimentally determined, the structure for Amy9 was predicted using AlphaFold2. The models of each CAZyme are colored by the domain annotation, similar to (a). For Amy4 and Amy10, the location of the active sites is outlined. **(d)** All structures retain the consensus alpha/beta-barrel-fold, centered around an 8-sheet beta barrel, and the catalytic triad consisting of two aspartate and a glutamate residue. Overlay of the four experimental models and the predicted model is shown. **(e)** A detailed look at the active site of Amy4. A maltotriose substrate from PDB 7DCG was docked into the active site, the catalytic residues are shown and labelled, and the enzyme subsite +1, which harbors the reducing-end glucose unit, is indicated. **(f)** In comparison, the active site of Amy10 accommodates the branched glucose chain in the same subsites as the linear chain in Amy4. At subsite +1, above the catalytic residues, space is required for the main polysaccharide chain, in this case maltopentaose, docked from Amy12, PDB 7LSA. The enzyme subsite +1 is indicated. **(g)** The active site of Amy4 (turquoise) overlaid on that of Amy10 (beige). The gate limiting Amy4 active site accessibility at the reducing end of the substrate, consists of the C-terminal loop of β5 with bulky residues His-353 and Trp-354. **(h)** Even though Amy4 and Amy16 have similar active sites, an additional large domain limits accessibility of the pocket to the non-reducing end of the polysaccharide substrate, here shown as an arrow.

To understand the role of each enzyme, we expressed and purified full-length Amy9, Amy10, Amy12, and Amy16, along with the truncated mutant Amy4Δdoc, which lacks the C-terminal dockerin domain, to assess enzyme activity while preventing self-aggregation of Amy4 through intramolecular cohesin-dockerin interactions. We used a thin-layer chromatography (TLC) assay to determine the preferred substrates of each enzyme. We tested the ability of each enzyme to hydrolyze the two most common glycosidic bonds occurring in RS: (1,4)- and (1,6)-glycosidic bonds. As model substrates for amylase activity, we used maltoheptaose (M7) and the more complex cornstarch (CS). As a model for pullulanase activity, we used pullulan. These experiments showed that Amy4Δdoc and Amy9 degraded only M7, while Amy16 degraded M7 and CS, with residual activity against pullulan. In contrast, Amy10 and Amy12 degraded pullulan, but neither M7 nor CS (Fig. 3b, Fig. S8a). Taking into account the individual enzyme activities and the selective enrichment of Amy4, Amy9, and Amy16 in the amylosome after growth on RS, we expected these enzymes to play key roles in RS degradation. A complementary relationship between Amy4 and Amy16 emerges, where Amy16’s broad specificity enables hydrolysis of multiple glycosidic bonds, while Amy4’s more defined specificity refines RS degradation, together ensuring efficient substrate breakdown and potentially compensating for the low abundance of a pullulanases in the proteome.

Next, we aimed to identify the structural features that would explain the substrate preferences of the key amylosome enzymes. The structure of the catalytic domain of Amy12 was previously determined^23^, revealing the active site of the protein and its mode of substrate binding. We used cryo-EM single-particle analysis to determine the structures of the catalytic domains of Amy4, Amy10, Amy12 and Amy16 at global resolutions of 2.8 Å, 3.1 Å, 3.1 Å and 4.6 Å, respectively. Here, we compare our atomic models for Amy4, Amy10 and Amy12, to a flexibly fitted predicted model for Amy16 and a predicted model for Amy9 (Fig. 3c). All proteins share the canonical structural features of GH13-family CAZymes: an alpha/beta barrel fold, consisting of 8 alpha-helices arranged around a central barrel consisting of 8 parallel beta-sheets, as well as the catalytic triad consisting of two aspartic acids and a glutamate, located at the active sites (Fig. 3d).

Given the observed similarities in the catalytic domain between the amylases Amy4, Amy9 and Amy16, and the two pullulanases Amy10 and Amy12, respectively, we set out to compare between Amy4 and Amy10, to derive the structural basis for their differences in substrate specificities. A cutaway view through the active site of Amy4 reveals that the catalytic residues are embedded in a deep pocket. Docking a maltotriose substrate from the *Weissella cibara* GH13-family glucosidase (catalytic residue RMSD 0.8 Å, PDB: 7DCG, ref^27^) revealed a tight fit of the substrate into the catalytic site. The three glucose residues of the maltotriose occupy the full catalytic pocket, with the catalytic residues positioned to cleave off the reducing-end glucose at position +1. The substrate is enclosed by protein on three sides. This has functional implications: as the +1 position can only accommodate a monosaccharide, and similarly Amy4 can therefore only act as an exoglycosidase on unbranched substrates (Fig. 3e). In comparison, the catalytic residues of Amy10 are located in a shallow cleft. Docking of the cleaved maltotriose/maltopentaose substrate from Amy12 (catalytic residue RMSD 1.0 Å, PDB: 7LSA, ref^23^) illustrates the fit of branched substrates into the catalytic site. Here, position +1 sits on the protein surface and can thus be part of the main polysaccharide chain at a branching point (Fig. 3f). The difference in accessibility of the +1 position between Amy4 and Amy10/12 is caused by the C-terminal loop of the central barrel β5. Whereas the extensions of β4 and β7, which carry the catalytic aspartate residues, run parallel in both enzymes, the C-terminal loop of β5 with its bulky residues H383 and W384 blocks the pocket of Amy4 around position +1; the equivalent loop in Amy10 folds back towards β4 (Fig. 3g). The glucose residues located towards the non-reducing end of the cleavage site are similarly placed in both enzymes.

While focusing on the +1 position accommodating the reducing end of hydrolyzed bond provides the structural basis for the diverging substrate preferences between the amylases and pullulanases, it does not explain the remarkable substrate promiscuity of Amy16 and the specificity of Amy4. Here, accessibility towards the non-reducing end of the substrate may be a key difference between Amy4 and Amy16. While the active sites of both enzymes sit in a deep pocket, Amy16 lacks several loops, which in Amy4 narrows the pocket towards the non-reducing end. The catalytic site is more accessible in Amy16, thus facilitating entry for longer-chain substrates (Fig. 3h). Even though Amy9 and Amy16 cluster together based on their sequence^22^, and are predicted to have similar folds (Fig. 3c), Amy16 has a wider substrate availability. Experimentally confirming the structure of Amy9 could help explain these differences.

In conclusion, the experimental structures of the amylosome CAZymes Amy4, Amy10, Amy12 and Amy16 explain the basis for the divergent substrate preferences between amylases and pullulanases and suggests a basis for complementarity of the specificity of Amy4 with the promiscuity of Amy16, potentially enabling simultaneous engagement to enhance RS degradation efficiency. Based on our functional and structural characterization of the amylosome enzymes, we hypothesize that this complementarity may play an important role for RS degradation.

### Synergistic effects between Amy4 and Amy16 as RS-degrading enzymes

To further elucidate the roles of Amy12, Amy10, Amy9, Amy4, and Amy16 in amylosome activity and to test our hypothesis of their complementary functions, we examined their activity against RS. Our results identify Amy4 and Amy16 as the primary enzymes driving RS decomposition and support our hypothesis of their complementary functions, as they demonstrate significant synergistic activity in this process.

In our experimental setup, the enzymes were incubated with HiMaize 958 RS, and product formation after 3 h was evaluated using TLC and the bicinchoninic acid (BCA) assay to quantify reducing sugars (Fig. 4a). The TLC indicated that only Amy4Δdoc and Amy16 can release oligosaccharides from RS (Fig. S8b), supported by reducing sugar quantification, where Amy4Δdoc released 0.4 ± 0.2 mM, while Amy16 released 1.1 ± 0.2 mM reducing sugars (1 µM enzyme, mean ± SD, 4 biological replicates, Fig. 4b). Contrary to our initial hypothesis, Amy9 did not release reducing sugars from the tested RS above background levels. However, its previously observed activity on a different RS type suggests a finely tuned enzymatic interplay that depends on the specific RS substrate^22^. The strong activity of Amy4 and Amy16 is consistent with our proteomic analysis, which showed that both enzymes were strongly enriched on RS. When all enzymes were incubated together, the released product reached 2.4 ± 0.2 mM, significantly exceeding the sum of individual enzyme activities (1.7 ± 0.2 mM, paired t-test, p = 0.01). This represents an increase of over 40%, highlighting a substantial and significant synergistic effect (Fig. 4c).

**Figure 4:**
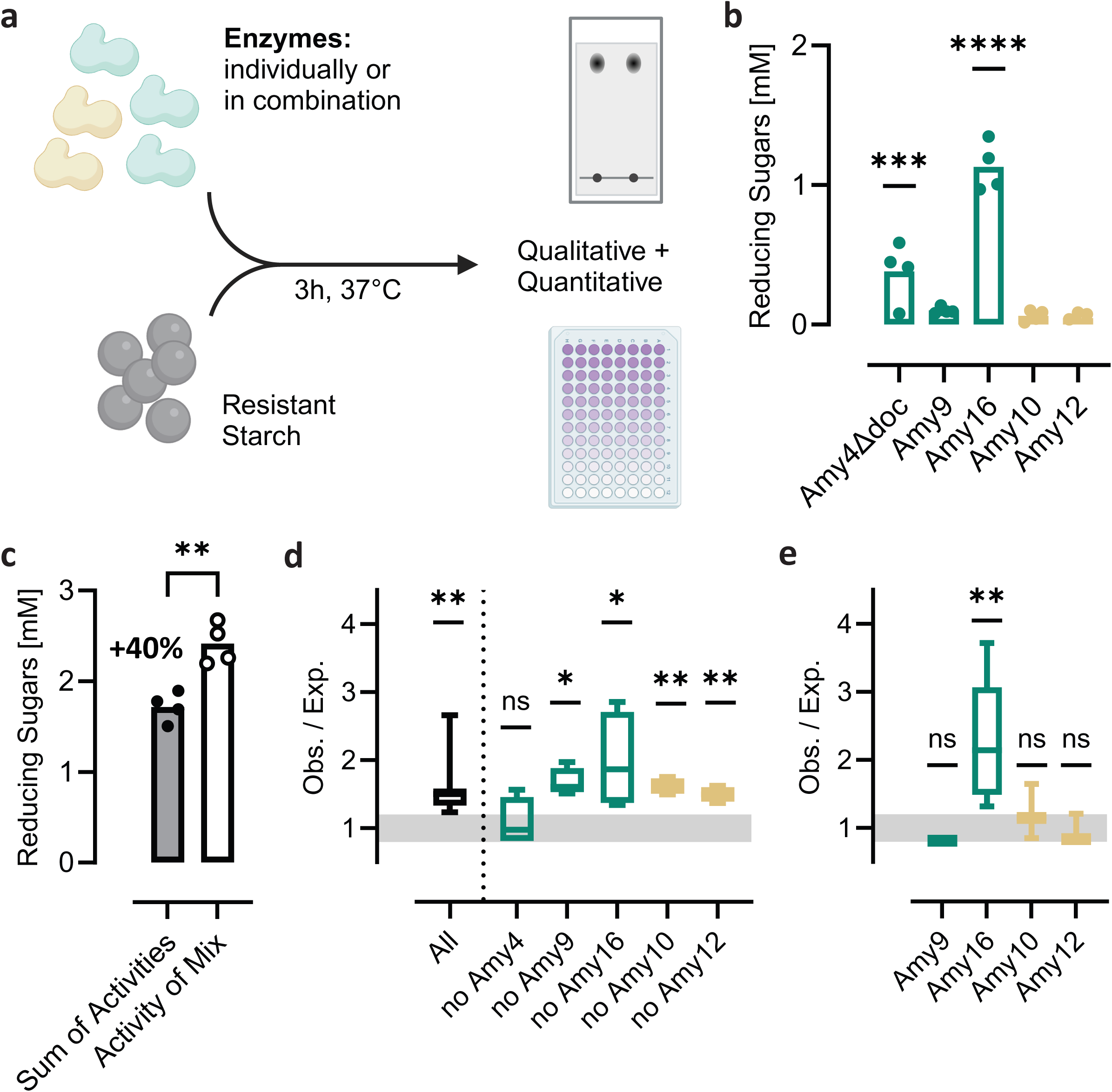
Synergistic effects between Amy4 and Amy16 as RS-degrading enzymes. **(a)** To test the RS-degradation capability of the amylosome CAZymes, the purified enzymes were incubated with resistant starch either individually or in combinations for 3 h at 37°C. After the reaction time, product formation was assessed qualitatively by thin-layer chromatography (Fig. S8b) and quantitatively using a BCA assay. **(b)** The production of reducing sugars quantified by BCA assay. Only Amy4Δdoc and Amy16 show significant reducing sugar production (one-way ANOVA with Dunnett’s multiple comparison test). **(c)** If all enzymes are incubated together, more substrate is produced than expected by the sum of activities (paired t-test). **(d)** Synergistic effects were quantified as the ratio of observed over expected product formation. In a leave-one-out series, no significant (ns) synergism was detected without Amy4Δdoc (90% CI shown, paired t-test, expected vs. observed product formation). **(e)** Pairwise synergism with Amy4Δdoc is only observed forAmy16, all other combinations not significant (ns). 90% confidence interval (CI) and results of paired t-test, expected vs. observed product formation, are shown. Panels b, c, d: 4 biological replicates, panel e: 3 technical replicates.

To further investigate this synergistic interaction among the five amylosome CAZymes, we quantified the individual contribution of each enzyme to the observed synergy, expressed as the ratio of observed over expected product formation. To this end, we measured the activity of enzyme combinations in which one CAZyme was omitted from the reaction mixture. Our results show that Amy4Δdoc is the key component driving synergism, as its absence resulted in the loss of significant synergistic effects. In contrast, the other enzymes were not essential for synergism to occur (Fig. 4d). When we tested whether synergistic effects could be captured in a minimal two-enzyme system by combining Amy4Δdoc with each of the other enzymes in the presence of RS, we found that Amy4Δdoc exhibited pairwise synergism exclusively with Amy16, but not with any other CAZymes (Fig. 4e). This finding not only corroborates our expression and structural predictions regarding their RS decomposition and complementary functions, but also extends them by demonstrating a direct synergistic interaction between the two enzymes.

### Cell-bound amylosome architecture enforces Amy4-Amy16 association for synergistic RS degradation

Based on our structural analysis and the potential complementarity of Amy4 and Amy16 in substrate interaction, we hypothesized that their physical proximity, maintained through a direct interaction, is important for their pronounced synergistic activity. This hypothesis was supported by the fact that wild type Amy4 contains both a cohesin and a dockerin domain (Fig. 3a), allowing it to function as a scaffoldin and interact with other amylosome proteins via its cohesin domain^19^.

To test the importance of the scaffoldin function, we compared the binding behavior and synergism of two truncated Amy4 mutants: Amy4Δdoc, lacking the dockerin domain, and Amy4Δcoh, lacking the cohesin domain. Both mutants retained catalytic activity against RS, as confirmed by TLC and reducing sugar quantification (Fig. S13).

Our results show that the Amy4 cohesin domain is essential for its interaction with Amy16 and for their synergistic activity. This was evident as Amy4Δdoc formed a stable complex with Amy16 during size exclusion chromatography (SEC) and showed significant synergistic effects in combination with all enzymes or solely with Amy16 (Fig. 5a). In contrast, Amy4Δcoh failed to bind to Amy16 and failed to exhibit significant synergistic effects (Fig. 5b). Taking together the binding, structural, and activity data, these findings indicate that Amy4 functions as a scaffoldin for Amy16, ensuring physical proximity, which significantly enhances *R. bromii’*s ability to degrade RS by facilitating synergistic CAZyme activity (Fig. 5c).

**Figure 5:**
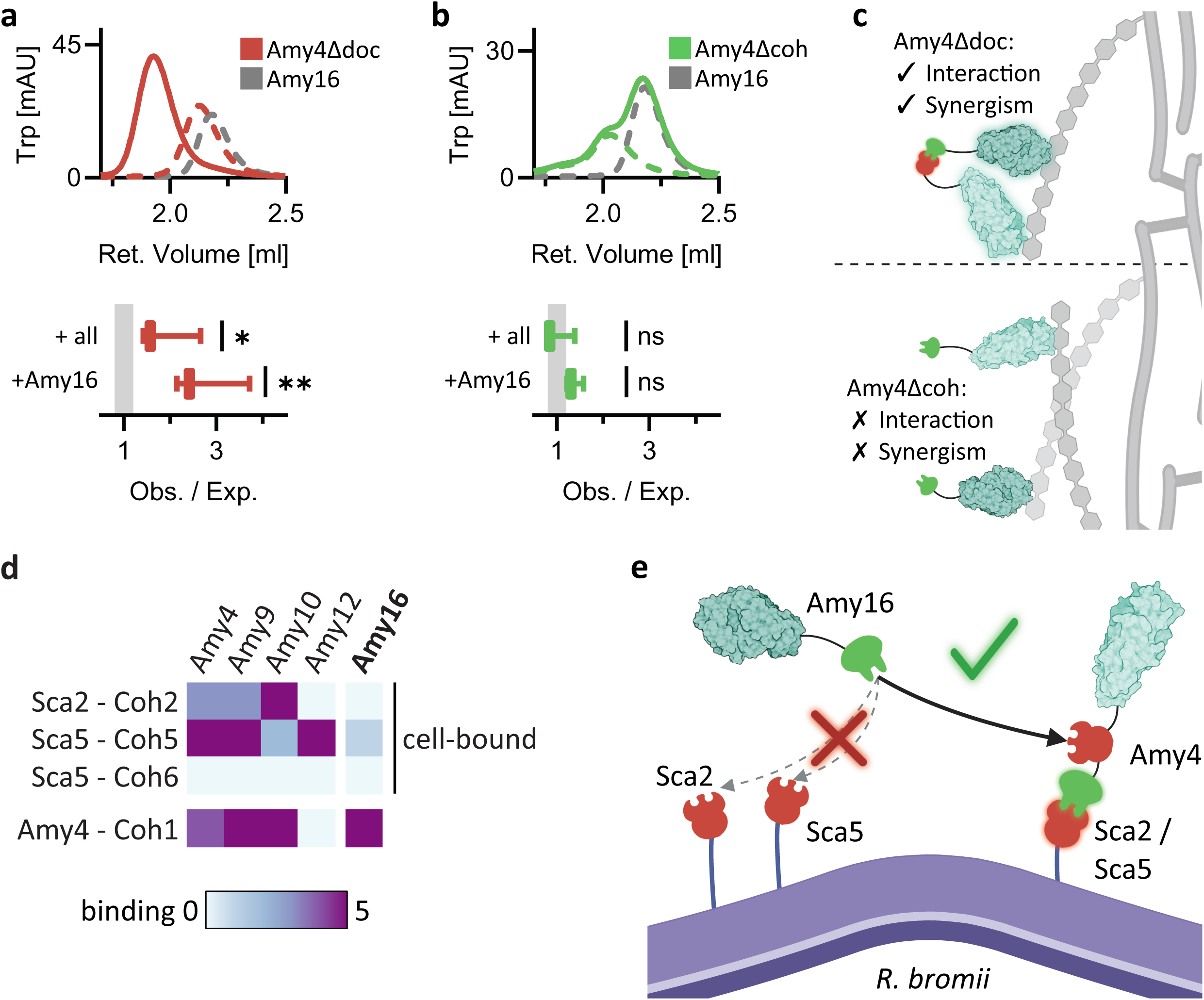
Cell-bound amylosome architecture enforces Amy4-Amy16 association for synergistic RS degradation. **(a)** Size-exclusion chromatography (SEC) profile showing tryptophan (Trp) fluorescence during elution of Amy4Δdoc, Amy16 (dashed), or their combination (continuous line). Amy4Δdoc elutes together with Amy16 in a single peak. Below, the observed over expected product formation of Amy4Δdoc incubated on RS either with all other amylosome CAZymes or with Amy16 alone. 90% confidence interval (CI) and results of unpaired t-test (expected vs. observed product formation) are shown. **(b)** SEC profile showing the Trp fluorescence during elution of Amy4Δcoh, Amy16 (dashed), or their combination (continuous line). The combination elutes in two separate peaks. Below, the observed over expected product formation of Amy4Δcoh incubated either with all other amylosome CAZymes or with Amy16 alone, indicates no significant (ns) synergistic effect. 90% CI and results of unpaired t-test are shown (expected vs. observed product formation). **(c)** A model linking complex formation and synergistic activity. Amy4Δdoc, which does interact with other enzymes, exhibits synergism. Amy4Δcoh, which does not interact, does not induce synergism. **(d)** Interaction data for the dockerin domains in the amylosome CAZymes with the cohesin domains in Amy4 or the cell-bound scaffoldins Sca2 and Sca5. Data from refs^19,22^, on the five-point scale from ref^19^. Amy16 does not bind to either cell-bound scaffoldin (Sca2 or Sca5), but binds strongly to Amy4. **(e)** The interaction data provides the basis for a model, in which all Amy16 in cell-bound amylosomes must be found in complex with Amy4, as there is no alternative mechanism for its recruitment to the cell. Only catalytic, cohesin and dockerin domains are shown in this schematic, for full model see Fig. S15. Activity data in panel a,b: 3 technical replicates

As our data suggest that the physical interaction between Amy4 and Amy16 enables their synergistic activity via cohesin-dockerin interactions, we further investigated scaffoldin interactions within the amylosome. Specifically, we examined how scaffoldins interact with the five amylosome CAZymes to better understand amylosome structure and function in RS degradation.

Remarkably, our results reveal that for Amy16 to associate with the cell wall-bound amylosome, it must be physically connected to Amy4 via cohesin-dockerin interactions, as it is the only amylosome CAZyme that lacks direct binding capability to cell-bound scaffoldins Sca2 and Sca5 (Fig. 5d). This suggests that compositional flexibility within the amylosome is constrained specifically for Amy16. Since Amy16 does not directly bind Sca2 or Sca5 but is highly abundant in the cell-associated proteome, we propose a model in which Amy4 acts as a cell-surface adapter for Amy16 (Fig. 5e). This model explains the strong enrichment of Amy4 and Amy16 (45 ± 3% and 14 ± 4% of amylosome proteins, respectively) in the cell-bound amylosome after growth on RS. In this organization, Amy4 is the sole recruiter of Amy16 to the cell surface, ensuring their spatial proximity and unlocking their synergistic effects in RS degradation. These findings suggest that amylosome architecture enforces close association of Amy4 and Amy16, positioning them as key players in amylosome function and RS breakdown.

### Integrative model of the amylosome highlights flexibility and efficiency

Next, we integrated our multi-scale data on the amylosome into a single model, encompassing population-level proteomics, nanometer-resolution cryo-ET and biochemical interaction studies. At the protein coordinates, which we derived from cryo-ET, we fit either Amy4, Amy9, Amy10, Amy12, Amy16 or “other” amylosome components, according to their occurrence in our proteomics dataset. We added CW and CM segmentations and mapped back the ribosomes from STA (Fig. 6a). When comparing the fructose- and RS-grown cells, the versatility of the amylosome is apparent: while the overall architecture and density is comparable between both conditions, the amylosomes after growth on RS are highly enriched in amylase enzymes (Video S1). After growth on fructose, in contrast, only a third of the proteins in the amylosome comprise CAZymes – the cell expresses many more accessory proteins (Video S2).

**Figure 6:**
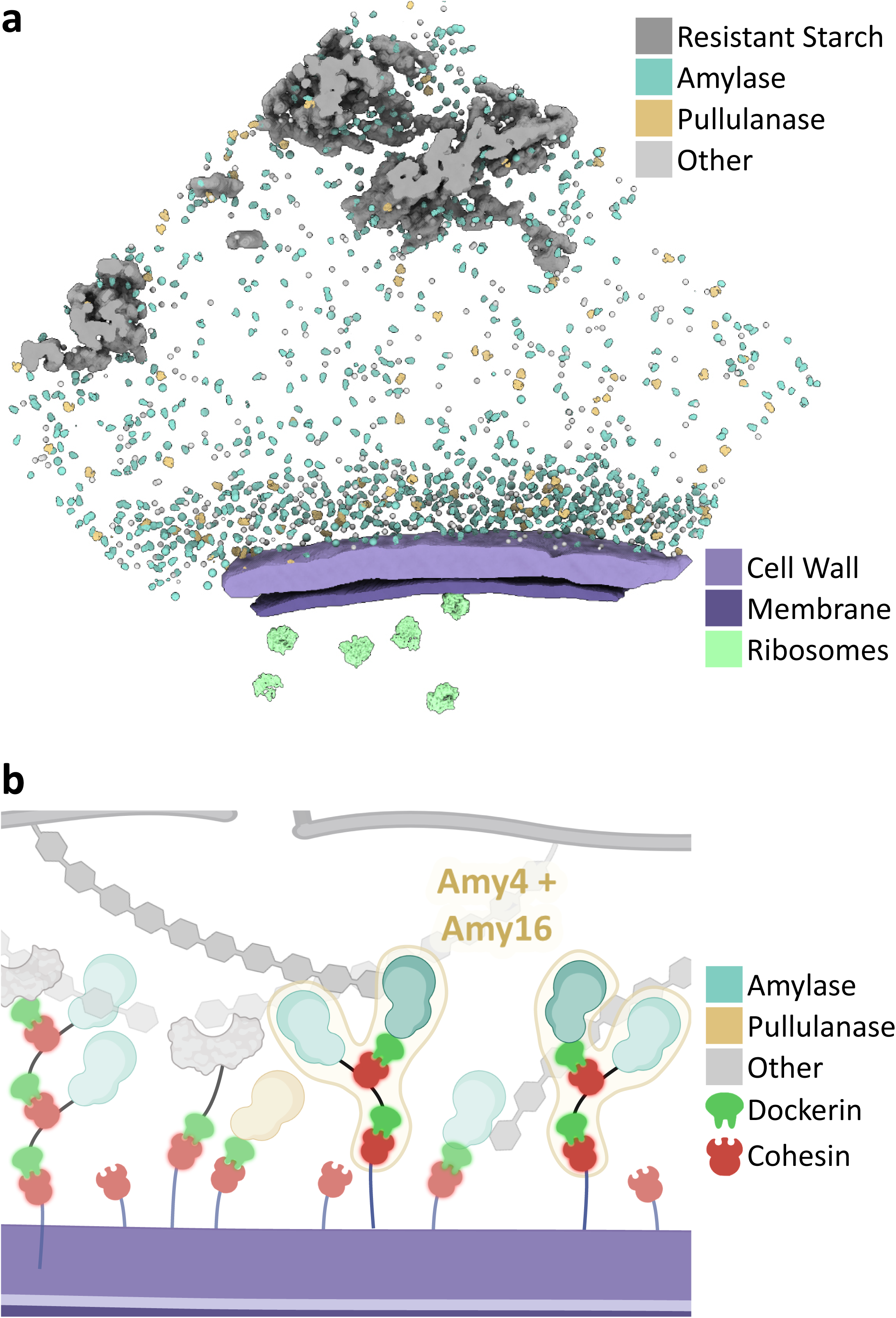
An integrative model of the amylosome highlights flexibility and efficiency. **(a)** An integrative model of the extracellular RS-degrading machinery of R. *bromii* grown on RS. On the basis of cryo-ET data, an integrative model containing cell wall, cell membrane, ribosomes and the extracellular proteins was built. This model highlights the versatility of the amylosome: After growth on RS, most of the amylosome is occupied by amylases. See also Videos S1 (RS) and S2 (fructose). **(b)** A schematic view of the amylosome fills in components which could not be modelled by cryo-ET, such as the scaffoldin proteins mediating amylosome adherence to the cell. After growth on RS, the synergistically active Amy4/Amy16-complex, which accounts for 30% of amylosome proteins.

The fact that overall amylosome architecture can be maintained, while the proteins it contains can be flexibly adapted can be directly traced to the modular architecture of the amylosome. The scaffoldins can remain in place, while the dockerin-containing proteins are flexibly exchanged. This is only possible due to the promiscuity of the cohesin-dockerin -interactions between the amylosome components (Fig. 5d, Fig. S14, Table S4). A schematic model allows further integration of the observed binding patterns and pronounced synergistic activity of Amy4 and Amy16 (Fig. 6b). During growth on fructose, when the enzymatic activity is not required, this complex accounts for only 10% of amylosome protein (Amy4 17 ± 1%, Amy16 5 ± 1%). During growth on RS, the complex accounts for 30% of amylosome proteins (Amy4 45 ± 3%, Amy16 14 ± 4%). Their high abundance and high affinity for each other enforce complex formation and together unlock the synergistic effects required to access this complex carbon source.

## Discussion

In this study, we present a comprehensive physiological, structural, and functional understanding of amylosome-mediated RS metabolism by *R. bromii*, a keystone species of the gut microbiome. We first leveraged cryo-ET to visualize the amylosome *in situ*, but could not observe any differences in appearance at this level. Notably, our analysis also did not detect amylosome densities in a cell-free form, suggesting that cell-bound amylosomes play a dominant role in RS degradation, consistent with previous studies^19^. We then generated a quantitative proteomics dataset with spatial and temporal resolution, highlighting the adaptability of the amylosome. Through biochemical assays, we identified Amy4 and Amy16 as synergistic partners for RS degradation, and identified their assembly in an amylosome-like fashion as a key to their synergism. Using combinatorics, we identified that Amy4 serves as an amylosome adapter to Amy16, thereby enforcing their assembly and suggesting synergism in the amylosome *in situ*.

Even though our comprehensive proteomics dataset uncovered precise and complex patterns of nutrient adaptation, the underlying regulatory mechanisms remain unknown. Indeed, unlike other starch-degrading bacteria in the gut microbiome, *R. bromii*’s CAZymes are not organized in polysaccharide utilization loci, but are instead dispersed throughout the genome^22^, suggesting that distinct regulon systems, potentially mediated by specific promoters and regulatory elements, control their expression.

Importantly, while the amylosome remains structurally stable, our findings reveal that its composition, combinatorics, and function are highly variable and responsive to different carbon sources. Within the amylosome, turnover could be facilitated by breaking down unnecessary amylase components, allowing newly synthesized proteins to be incorporated^28^. Potentially, the dockerin-containing protease Doc17 could play this role, as it strongly contributes to the differentiation of the “generalist” amylosomes after growth on fructose or starch, but not in those adapted for pullulan or RS in a PCA. This adaptability is further illustrated in the updated models for cell-associated amylosomes grown on all four carbon sources, as shown in Figure S15.

On a structural level, our characterization of Amy4, Amy10, Amy12, and Amy16 provides the molecular basis for their distinct substrate specificities. Pullulanases Amy10 and Amy12, which accommodate branched substrates, likely exhibit lower affinity for linear substrates. In contrast, Amy4 possesses a deep catalytic pocket, allowing precise coordination of linear substrates. This study contributes to a growing body of structural research on GH13-family CAZymes, which share a conserved fold yet exhibit diverse substrate preferences and are abundant in the human gut microbiome^29^. Biochemical assays and interaction studies identified Amy4 and Amy16 as the principal RS-degrading enzymes, exhibiting strong synergistic activity critically dependent on their spatial organization. The biochemical basis for this synergism between structurally similar enzymes when they are assembled into an amylosome remains elusive. Potentially, the complementarity between the promiscuous Amy16 and the specific Amy4 could permit simultaneous activity on the same polysaccharide chain, or the increased number of CBM domains in the complex improves substrate-binding through avidity effects, as has been observed for other starch-degrading enzymes with multiple CBMs^30^.

While the amylosome displays high combinatorial flexibility, we show that combinatoric constraints enforce the close proximity of Amy4 and Amy16 on the cell-associated amylosome, ensuring efficient RS breakdown. For example, in response to growth on RS, the enrichment of Amy4, which interacts with a wide array of amylosome components and is otherwise subject to competition within the complex, increases its availability across the amylosome. This reduces competition for its cohesin domain and favors its interaction with Amy16, which also represents a large proportion of the amylosome components. This co-enrichment enhances the likelihood of Amy16 docking onto Amy4’s cohesin domain, prioritizing their interaction over other CAZymes that would otherwise compete for the same binding site. This selective recruitment maximizes their synergistic RS degradation in both cell-bound and cell-free fractions. Such proportional modulation likely enables distinct combinatorial assemblies that optimize enzymatic function based on substrate availability across different conditions.

Taken together, this work provides key insights into amylosome organization, enzyme interactions, and RS degradation, highlighting the molecular basis for *R. bromii’*s role as a keystone species in the gut microbiome. Additionally, our study delivers valuable physiological, biochemical, and structural resources to the scientific community, offering a foundation for future research into *R. bromii* function and its ecological significance. The synergy and combinatorics - driven mechanism for efficient RS degradation identified in this study may help in the future design of better pre- and probiotics, which might restore RS-degrading ability to all microbiomes, counteracting the dysbiosis frequently induced by an industrialized diet.

## Methods

### Bacterial culture

*Ruminococcus bromii* L2-63 was grown in Hungate tubes in 8 mL of M2 anaerobic media^31^, supplemented with 0.2% of either fructose, pullulan (Merck), starch (from potato, Merck) or resistant starch Hi-Maize 958 (Ingredion, kindly provided by Prof. Harry Flint, University of Aberdeen) and incubated at 37°C. Growth curves were measured on each carbon source to determine mid-logarithmic, end-logarithmic and late-stationary phases (Fig. S3). Optical densities were measured for fructose, pullulan, and starch while quantitative PCR was used to determined 16S rRNA copy numbers per µL in resistant starch cultures using 16S RNA universal primers covering the V2 and V3 regions of the gene (5′-TTTGATCNTGGCTCAG-3′ and 5′-GTNTTACNGCGGCKGCTG-3′) as described previously^32^.

### Proteomics sample preparation

Cultures were grown as biological triplicates on all four carbon sources and harvested at three time points (mid-logarithmic, end-logarithmic and late-stationary phases) by centrifugation at 5,000 g for 10 min. The pellets were treated using the shaving approach as described previously^33^. Supernatant samples were frozen, stored at −80°C and sent to University of Greifswald for further preparation. A volume of 1 mL of the supernatant fluids were used for protein enrichment with 20 µg StrataClean^TM^ beads as previously described^34^. The dried StrataClean^TM^ beads were resuspended in 30 µL 2x reducing Laemmli sample loading buffer, incubated for 10 min at 98°C, and loaded on an SDS gel. Proteins were separated via SDS-PAGE and in-gel digested as previously described^35^ and desalted using U-C18 Zip Tips (Merck Millipore) according to the manufacturer’s instructions.

### LC-MS/MS measurement and data analysis

Tryptic peptides were separated on liquid chromatography system (EASY nLC) with an in-house built 20 cm column (inner diameter 100 µm, outer diameter 360 µm) filled with ReproSil-Pur 120 C18-AQ reversed-phase material (3 µm particles, Dr. Maisch GmbH, Ammerbuch, Germany). The peptides were loaded with solvent A (water MS grade in 0.1% acetic acid (v/v)) and afterwards eluted within a non-linear 100 min gradient ranging from 5% to 99% solvent B (99.9% acetonitrile (v/v), 0.1% acetic acid (v/v)) at the constant flow rate of 300 nL/min. Eluting peptides were injected into an LTQ Orbitrap XL mass spectrometer (data-dependent acquisition mode). Full scans were recorded, and most abundant five precursor ions were selected for fragmentation (full scan resolution 30,000; mass range 300 to 2,000 m/z).

MS *.raw data were searched with MaxQuant^36,37^ (version 2.0.3.0) and its implemented search engine Andromeda^38^ against a protein database obtained from Genbank assembly GCA_900291485.1 (2,111 proteins) and MaxQuant’s generic contamination list. The following parameters were used: digestion mode, trypsin/P with up to two missed cleavages; variable modification methionine oxidation, protein N-termini acetylation, and maximal number of 5 modifications per peptide; activated label-free quantification (LFQ) option with minimal ratio count of two and “match-between-runs” feature for biological replicates and classical normalization for cell-surface proteome. Proteins were identified with at least two unique peptides; peptide-spectrum match and protein false discovery rate were set to 0.01.

For data analysis the software Perseus^39^ (version 2.0.3.0) was used. The data from MaxQuant output files were filtered for contaminations, reverse entries and only identified by site hits. Since classical normalization for the secretome data set failed, the “LFQ intensities” were Z-score normalized. A protein was only quantified with “LFQ intensities” if it was present in at least two of three replicates. The quantified abundances for all replicates for cell-associated and supernatant proteome are listed in Tables S1 and S2, respectively.

All entries in the strain-specific database were additionally annotated for CAZymes using the dbCAN3 web server^40^ (results filtered to domains found by at least 2 tools) and the cellular localization was annotated using the pSortB 3.0 web server^41^. We considered proteins as putative amylosome components if they had an annotated dockerin or cohesin domain or had a predicted cell wall localization. For PCA, missing values were imputed based on the probabilistic minimum method.

### Cloning and constructs

Primers for the design of Amy4, Amy9, Amy10, Amy12, Amy16 and dockerin-containing constructs in pET28a are listed in Table S3, and used with *R. bromii* L2-63 genomic DNA as a template for the PCR. Plasmid GFP-Doc13a was previously described^22^. Residue numbering used throughout this paper is in reference to NCBI GenBank sequences, as indicated in Table S4.

To prepare the Amy4Δdoc and Amy4Δcoh mutants, the wildtype plasmid was linearized by PCR with primers omitting the deletion and an overhang on the forward primer (Table S3). The dockerin deletion removes Leu-1294–C-terminus, the cohesin deletion removes Ser-1138–Asp-1265. The template was removed from the reaction mix by DpnI digest for 3 h at 37°C. *E. coli* DH5a cells were transformed with the linearized vector and allowed to recover for 3 h before plating. The next day, several clones were selected for sequencing to obtain the desired plasmid.

### Protein expression and purification

Purifications of enzymes and dockerin-containing proteins were performed as follows: *E. coli* BL21* cells were transformed with the desired plasmid and plated. A single clone was extracted and used to inoculate 3 x 0.5 L pre-warmed Luria-Bertani (LB) medium, incubated at 37°C with shaking (120 rpm). At OD ∼ 1, 0.25 L cold LB medium and isopropyl ß-D-1-thiogalactopyranoside (IPTG) to a final concentration of 1 mM were added, and the flask was transferred to a pre-cooled incubator for overnight expression at 16°C with shaking (90 rpm). Cells were harvested by centrifugation at 5,000 x g for 20 min at 4°C. Pellets were washed with cold phosphate-buffered saline (PBS, 12 mM phosphate, 137 mM NaCl, 2.7 mM KCl) and harvested again for 15 min at 5,000 x g, 4°C. After removing the supernatant fluids, pellets were flash-frozen and stored at −80°C until purification.

GFP-Doc13a was expressed as follows: Transformed *E. coli* BL21 DE3 cells were grown in 3 x 0.5 L LB supplemented with 2 mM CaCl_2_. At OD ∼1, they were induced with 3 mM IPTG and transferred to a pre-cooled incubator for overnight expression at 18°C, with shaking (110 rpm). Cells were harvested by centrifugation for 15 min at 6,000 x g, 4°C. Pellets were washed with cold PBS and harvested again by centrifugation for 15 min at 6,000 x g, 4°C.

For purification, pellets were thawed on ice and incubated with lysis buffer (25 mM Tris, 137 mM NaCl, 2.7 mM KCl, 5 mM imidazole, 2 mM phenylmethylsulphonyl fluoride (PMSF), cOmplete protease inhibitors (Roche), pH 7.4 + 10 mM CaCl_2_ for enzymes).

Cells were additionally lysed by sonication, and debris removed by centrifugation at 20,000 x g, 1 h, 4°C. The supernatant fluids were syringe-filtered to 0.45 µm and loaded on a HisTrap FF column (Cytiva). After a wash of 10-15 column volumes (CV) with 20 mM imidazole, the proteins were eluted using a gradient up to 500 mM imidazole while collecting 1 mL fractions. The fractions containing the proteins were dialyzed into Tris-buffered saline (TBS) + 10 mM CaCl_2_, pH 7.4 and concentrated using a spin concentrator (enzymes) or directly concentrated without dialysis (GFP-Doc13a), before injecting 500 µL on a Superdex 200 increase 10/300 GL increase size exclusion column (Cytiva), run at 0.3 mL/min for 1.5 CV with SEC buffer (TBS + 10 mM CaCl_2_ + 2 mM dithiothreitol (DTT), pH 7.4). Fractions containing the proteins of interest were pooled and concentrated using spin concentrators.

While Amy9, Amy10, Amy12 and Amy16 were eluted as a monodispersed peak during SEC, Amy4 showed significant signs of aggregation. We presumed that this was caused by homo-oligomer formation through intermolecular cohesin-dockerin interactions. We therefore cloned and purified AmyΔdoc, a mutant without the C-terminal dockerin domain, which did not show signs of aggregation. The activity profile of this mutant matched the wild-type enzyme, and it was thus used for further characterization.

Enzymes were used directly after SEC purification or flash-frozen and stored at −80°C. GFP-Doc13a was diluted to a final concentration of 0.25 mg/mL in 1:1 glycerol:SEC buffer, flash-frozen and stored at −20°C.

### GFP-Doc13a staining and light microscopy

To visualize the cohesins exposed on the cell-surface, cells were allowed to interact with GFP-Doc13a. Bacteria were cultured on the appropriate carbon source as described above. A 1-mL volume of culture was collected, and the cells were harvested at 5,000 x g for 5 min. Cells were resuspended in 200 µL PBS and washed 3 times (pellet 3 min at 5,000 x g, resuspend). The final resuspension was done in 160 µL PBS + 10 mM CaCl_2_ + 40 µL (10 µg) GFP-Doc13a + 1:1,000 SPY650-DNA (Spirochrome). Cells were incubated for 1 h at room temperature on a rotating platform. After cohesin-dockerin binding, the cells were again washed 3 times. A volume of 10 µL of the cell suspensions was added on a glass slide and air-dried in the dark for 1 h before adding DAKO mounting medium (Agilent) and closing with a cover slip, sealed with nail polish. Samples were imaged on an Olympus IX83 widefield microscope using a 100x NA 1.4 oil objective with a z-step of 280 nm, using brightfield, Cy5 and Alexa488 filter sets. Image stacks were bleach-corrected and converted to tiff format using vsi2tif (https://github.com/bwmr/vsi2tif) before deconvolution in SVI Huygens (Scientific Volume Imaging, Hilversum, Netherlands).

### Confocal microscopy of cell-associated scaffoldins

To validate the presence of the cell-associated scaffoldins Sca2 and Sca5 on the cell surface, cells grown on fructose were immuno-labelled and processed as described previously^20^ using specific chicken antibodies (Aluma-Bio, Israel) against full-length Sca2 (at 1:1,000 dilution) or the first cohesin of Sca5 (at 1:250 dilution). Alexa Fluor® 647 goat anti-Chicken IgG (ThermoFisher Scientific) was used as secondary antibody at 1:500 dilution, and cells were imaged using a 3i Marianas spinning disk confocal microscope equipped with a Yokogawa W1 module and Prime 95B sCMOS camera. Excitation laser (absorbance 637 nm and emission filter 672–712 nm) was achieved using a Å∼100 Zeiss Plan-Apochromat 1.4 NA DIC oil objective.

### Affinity-based ELISA analysis of dockerin cohesin binding

The previously published protocol was followed^42^.

### Cryo-EM sample preparation

Purified proteins were concentrated to 0.3 mg/mL (Amy4Δdoc), 0.08 mg/mL (Amy10/12) or 0.1 mg/mL (Amy16). Au 200-mesh R1.2/1.3 holy carbon grids (Quantifoil) were plasma-cleaned for 35 s on “High” and 10 s on “Low” (Harrick Plasma PDC-32G). Plunge-freezing was conducted on a Vitrobot Mk. IV (FEI), the camber set to 4°C and 100% humidity. 3 µL of protein solution were applied to each grid, which was then blotted 3.5 - 7.5 s (blot force −1), before being vitrified using liquid ethane. Grids were stored in liquid nitrogen until imaging.

### Cryo-EM SPA data acquisition and analysis

Samples on grids were clipped into AutoGrids and loaded into Titan Krios G3i cryo-TEM (ThermoFisher Scientific) operated at 300 kV equipped with an energy filter (zero-loss mode, 20 eV width) and K3 direct electron detector (Gatan). Data acquisition was controlled using EPU, full acquisition parameters for each dataset can be found in Table S5.

Cryo-EM SPA processing was performed using cryoSPARC (Structura Biotechnology). Briefly, movies were preprocessed with patch-based motion correction and CTF estimation, before particle picking, 2D classification and refinement against *ab-initio* generated templates. The processing pipelines for each structure are listed in Figures S9 - S12. For Amy10, the 3DFlex workflow^43^ was used to retrieve additional density belonging to the accessory domains.

### Model-building

The models for Amy4 and Amy10 were built based on the AlphaFold2 predicted models^44^. Predicted models were docked into the densities using ChimeraX^45^, preprocessed using Phenix^46^. The models were then adapted in an iterative workflow with building in ISOLDE^47^ and real-space refinement in Phenix. The model for Amy12 was built based on the previously published^23^ x-ray structure (PDB: 7LSA), which was docked using ChimeraX and subjected to real-space-refinement in Phenix. Complete refinement and validation statistics can be found in Table S5.

### Enzyme activity assays

Enzymes were incubated at 1 µM final concentration in SEC buffer (as described above) with 5 mM maltoheptaose (Cayman Chem), 0.6% (w/v) pullulan (Sigma-Aldrich), 0.3% (w/v) cornstarch (Maizena), or 0.3% HiMaize 958 resistant starch (Ingredion) at 37°C for 1 h (M7, pullulan, CS) or 3 h while shaking at 300 rpm (RS). For each digestion, a substrate-only and a protein-only control was included.

### Thin-layer chromatography

Product formation was observed using TLC on 10 x 20 cm silica plates (Macherey-Nagel), following a previously published protocol^48^. On each plate, 1 µL of 10 mM glucose solution was spotted as a control. For digestions of M7, pullulan or CS, 2 x 1 µL of reaction mix were spotted, for RS degradation 3 x 1 µL. The TLC was then run in a pre-equilibrated chamber with an 85:20:50:50 acetonitrile : ethyl acetate : isopropanol : ddH_2_O eluent until the solvent front reached 3 cm below the top of the plate. After the solvent had evaporated, the plate was stained by spraying with 0.8% (w/v) N-(1-naphtly)ethylenediamine (Sigma-Aldrich) in 5% (v/v) sulfuric acid in methanol. After allowing the staining solution to dry, the plate was heated in an oven to 120°C for 10 min, until spots could be seen, and imaged.

### BCA assay

Reducing sugars were quantified using the bicinchoninic acid (BCA) assay. Briefly, 2 µL of reaction mixture or glucose standards were diluted in 60 µL ddH_2_O. These were incubated with 60 µL BCA solution prepared as described here^49^ at 80°C for 20 min. Then, 100 µL of each sample were transferred to a 96-well plate, in which absorption was measured at 562 nm.

Absorption values were corrected for background using a BCA + ddH_2_O control. Then, per-plate linear fit was calculated using the glucose standards, and unknown values interpolated. Concentrations of reducing sugars were corrected by a substrate-only control before plotting or further calculations.

To evaluate whether proteins acted synergistically, the expected and measured product concentrations were compared. To calculate the expected product formation, the individual activities of all proteins in a given set were summed, and compared to the product formed by the set of proteins together. Expected and measured activities were always compared within the same replicate, to minimize batch effects, using the paired t-test. For plotting of synergistic effects, the ratio of observed product formation over expected product formation for each replicate was calculated, with values > 1 indicating synergistic effects.

GraphPad Prism 10.3 (GraphPad Software, Boston, Massachusetts USA) was used for calculations and statistical analysis.

### Elution shift assay

Amy4Δdoc, Amy4Δcoh and Amy16 were purified from flash-frozen dialysate by SEC, using a HEPES [(4-(2-hydroxyethyl)-1-piperazineethanesulfonic acid)]-based buffer (25 mM HEPES, 137 mM NaCl, 2.7 mM KCl, 10 mM CaCl_2_, 2 mM DTT). Five samples were prepared for the analytical SEC run: Amy16, Amy4Δdoc and Amy4Δcoh individually, Amy16 + Amy4Δdoc and Amy16 + Amy4Δcoh. Each enzyme was used at 1 µM concentration. Samples were filtered using a spin column. 50 µL of each sample were applied to a Superose 6 increase 5/150 GL column, run at 0.2 mL/min, while measuring absorption at 280 nm and tryptophane fluorescence at 330 nm (Agilent 1200 Series HPLC).

### Cryo-ET sample preparation

*R. bromii* cells were grown for 20 h in M2 medium containing either fructose or RS as the sole carbon source. For fructose samples, cells were harvested for 5 min at 4,000 x g and resuspended in PBS to a final OD (600 nm) of 5. Cu 200-mesh R0.6/1 (Quantifoil) grids were plasma-cleaned for 1 min (Pelco EasyGlow). 3 µL of cell suspension were added, and grids were blotted from the back for 12 s before plunging into liquid ethane using a manual plunge-freezer.

For RS samples, suspension culture was diluted 1:1 with PBS and 3 µL were directly applied to glow-discharged Cu 200-mesh R2/1 grids (Quantifoil), and blotted from the back for 8 - 10 s before plunge-freezing in liquid ethane.

### Cryo-FIB milling

EM grids were clipped into AutoGrids with a cutout for FIB milling (ThermoFisher Scientific). FIB milling was conducted using a Gallium source operated at 30 kV either on a Auriga 40 CrossBeam (Zeiss) at the Centre for Microscopy and Image Analysis at the University of Zurich or an Aquilos 2 (ThermoFisher Scientific) at the EMBL Imaging Center in Heidelberg. In all cases, micro expansion joints^50^ were used.

At the Auriga, samples were sputtered with 6 nm of platinum before loading into the microscope, where they were additionally coated with an organometallic platinum layer. Sample thinning was controlled manually through the Nanopatterning and Visualization Engine, with current steps 240 pA, 120 pA, 50 pA and a final polishing step at 20 pA to a set thickness of 170 nm.

On the Aquilos 2, samples were sputtered with platinum directly after loading into the microscope and after polishing was complete. Additionally, a layer of organometallic platinum was added before FIB milling. Milling was controlled using AutoTEM (ThermoFisher Scientific), with current steps 1 nA, 0.5 nA, 0.3 nA and a two-step polishing with 50 pA and 30 pA to a set thickness of 150 nm.

### Cryo-ET data acquisition and preprocessing

Cryo-ET data of FIB lamellae were acquired using a Titan Krios (FEI) operated at 300 kV, equipped with an energy filter and K2 summit detector (Gatan). Acquisition was controlled using SerialEM 4.1^51^. First, grid overviews were acquired at 175x magnification to identify lamella sites. Then, a medium-magnification montage of each lamella was acquired at 6,500x magnification (calibrated pixel size 2.20 nm/px) to select positions for cryo-ET data acquisition using PACEtomo^52^. Low-magnification tomograms were acquired at 8,700x magnification (pixel size 1.67 nm/px) using a total dose of 20 e^−^/Å^2^ in a bidirectional acquisition scheme of 3° increments. Other tomograms were acquired at 33,000x or 105,000x magnification (calibrated pixel size 0.4039 nm/px and 0.1344 nm/px, respectively) in a dose-symmetric scheme with 3° increments starting from the lamella pre-tilt, with a total dose of 120-140 e^−^/Å^2^.

Frame series were saved as non-gaincorrected tif files and processed using the tomotools workflow (https://github.com/tomotools/tomotools): Frame series were gain- and defect-corrected and aligned into tilt images with MotionCor2^53^ correcting only for global motion, and then assembled into stacks using newstack from the imod package^54^. Then, tilts to be excluded from the tilt series due to obstructed or shifted field of view were determined manually, and the cleaned stacks were passed to AreTomo^55^ for tilt alignment. In case the tilt alignment was of low quality, alignments were re-done using etomo patch tracking or fiducal markers, if possible. In all cases, the aligned stacks were then dose-filtered using mtffilter and back-projected using tilt, both from the imod package. For figure preparation, additional tomograms were reconstructed from the even and odd frames, respectively, and denoised and wedge-corrected using cryoCARE^56^ or DeepDeWedge^57^.

### Visualization and segmentations

For quantification of amylosome density, only a subset of tomograms showing a clear side-view of CW and CM was used (n = 10 for RS, n = 11 for fructose).

Tomograms of the even and odd frames were reconstructed at bin 8 with a SIRT-like filter of 20 and used to train one DeepDeWedge model per condition. On the DeepDeWedge corrected tomograms, membrane and cell wall were segmented automatically using membrain-seg^58^ and cleaned using seg-select (https://github.com/bwmr/seg-select). RS was annotated manually in napari every 5 slices, and the annotations were interpolated and adapted to the density using custom Python scripts.

Putative amylosome components were picked on tomogram reconstructions at bin 4, SIRT-like filter 20, using sizepicker (www.github.com/bwmr/sizepicker), following a previously published approach^59^. Cytoplasmic picks were manually removed using napari_boxmanager (https://github.com/MPI-Dortmund/napari-boxmanager).

### Sub-tomogram averaging of ribosomes

Ribosome coordinates were obtained by template matching using pytom-match-pick^60^ on 3D-CTF corrected tomograms at bin 8, using a rescaled version of the *T. kivui* ribosome (EMD-16451) as a template.

For STA of ribosomes, tilt images and tilt alignment information were exported to WarpTools^61^ using the tomotools imod2warp and aretomo2warp commands. After CTF estimation for tilt images and tomograms, particle series in the Relion 5 convention^62^ were reconstructed at bin 4. After an initial alignment against a low-pass version of the *T. kivui* ribosome template, one round of 3D classification without alignment and another refinement, tilt series alignments and particle poses were refined using M^63^ (Fig. S2).

### Integrative amylosome model

The putative amylosome coordinates for each tomogram were split into random subsets representing Amy4, Amy9, Amy10, Amy12, Amy16 or “other” amylosome components. The sizes of these subsets were sampled from the distributions observed by quantitative proteomics. Then, the coordinates were assigned random angles and written out as .star files for rendering. All renderings were created in ChimeraX 1.8 using the ArtiaX plugin^64^.

### Illustrations

Illustrations in Fig. 4a, Fig. 5c/d and Fig. 6b were created using biorender.com and are available under the following URLs: https://biorender.com/c49f659 (Fig. 4a), https://biorender.com/4nx8bk0 (Fig. 5c), https://biorender.com/y31q9fi (Fig. 5e), https://biorender.com/3f1dh58 (Fig. 6b).

## Supporting information

Document S1, with Figs. S1 - S15

## Data and Code Availability

The mass spectrometry proteomics data have been deposited to the ProteomeXchange Consortium via the PRIDE partner repository^65^ with the dataset identifier PXD060149 (Project accession: PXD060149, Token: JW74CYXGTpuc). The proteomics data analysis pipeline, including annotated sequence database and the MaxQuant output files are available at Zenodo, doi: 10.5281/zenodo.15114156

The cryo-EM density maps and their associated models were deposited at EMDB and PDB, respectively: Amy4Δdoc (EMDB: 53097 / PDB: 9QF3), Amy10 (EMDB: 53101 / PDB: 9QF8), Amy12 (EMDB: 53103 / PDB: 9QFA), Amy16 (EMDB: 53102, PDB: 9QF9). The STA map of the *R. bromii* ribosome was deposited at EMDB, accession code EMDB: 53104.

## Author Contributions

Conceptualization, I.M., O.M.. Investigation: B.H.W., S.M., O.T.H., A.T.S., P.T., S.S., M.L., I.A., O.T.H., M.T.. Resources: S.M.. Data curation: B.H.W., S.M., A.T.S., L.L.. Writing – original draft: B.H.W. and S.M.. Writing – review & editing: B.H.W., S.M., A.T.S., E.A.B., O.M., I.M.. Visualization: B.H.W. and S.M.. Supervision: O.M., I.M., D.B.. Funding acquisition: O.M., I.M.

## Acknowledgements

This work was funded by grants from the German-Israeli Project Cooperation (DIP 2476/2-1) to I.M., the European Research Council (ERC 866530) to I.M., the Israel Science Foundation**-** Swiss National Science Foundation (ISF 1057/24 / SNSF 10000885) to I.M. and O.M, and the Novartis Foundation for Medical-Biological Research to O.M. and B.H.W.. Access to the cryo-FIB-SEM at the EMBL-IC was funded through iNEXT-Discovery (Proposal 24271) and facilitated on-site by Dr. Zhengyi Yang. The authors thank Prof. Harry Flint (University of Aberdeen) for providing Hi-Maize 958, Eva Setter-Lamed for technical help with enzymatic purification and Melanie Arndt (University of Zurich) for help with the elution shift assay. We also thank Sebastian Grund (University of Greifswald) for technical assistance in handling the proteomics samples and the mass spectrometric measurements. Cryo-EM/-ET imaging and FIB milling at the Zeiss Auriga were performed using equipment maintained by the Center for Microscopy and Image Analysis at the University of Zurich. Image acquisition with the spinning disk confocal microscope was performed in the Ilse Katz Institute for Nanoscale Science and Technology Shared Resource Facility under the direction of U. Hadad.

## Supplemental Files

Document S1: Figures S1 – S15.

Table S1: Excel table with all quantified protein abundances in the cell-associated proteome, log2-transformed LFQ values, per replicate. Relates to Fig. 2.

Table S2: Excel table with all quantified protein abundances in supernatant, z-score normalized, per replicate. Relates to Fig. 2.

Table S3: Primer sequences used in this study, relates to methods section.

Table S4: Sequence identifiers, activity and binding information for amylosome components and related free-enzymes. Relates to Fig. 2 and Fig. S5.

Table S5: Refinement and validation statistics for cryo-EM structure determination, relates to Fig. 3 and methods section.

Video S1: Rendering of an integrative model of the amylosome after growth on RS, relates to Fig. 6.

Video S2: Rendering of an integrative model of the amylosome after growth on fructose, relates to Fig. 6.

## References

1. Makki, K., Deehan, E.C., Walter, J., and Backhed, F. (2018). The Impact of Dietary Fiber on Gut Microbiota in Host Health and Disease. Cell Host Microbe 23, 705–715. 10.1016/j.chom.2018.05.012.

2. Spivak, I., Fluhr, L., and Elinav, E. (2022). Local and systemic effects of microbiome-derived metabolites. EMBO Rep 23, e55664. 10.15252/embr.202255664.

3. Flint, H.J., Scott, K.P., Duncan, S.H., Louis, P., and Forano, E. (2012). Microbial degradation of complex carbohydrates in the gut. Gut Microbes 3, 289–306. 10.4161/gmic.19897.

4. Koh, A., De Vadder, F., Kovatcheva-Datchary, P., and Bäckhed, F. (2016). From Dietary Fiber to Host Physiology: Short-Chain Fatty Acids as Key Bacterial Metabolites. Cell 165, 1332–1345. 10.1016/j.cell.2016.05.041.

5. Robertson, M.D., Bickerton, A.S., Dennis, A.L., Vidal, H., and Frayn, K.N. (2005). Insulin-sensitizing effects of dietary resistant starch and effects on skeletal muscle and adipose tissue metabolism. Am J Clin Nutr 82, 559–567. 10.1093/ajcn.82.3.559.

6. Le Leu, R.K., Hu, Y., Brown, I.L., and Young, G.P. (2009). Effect of high amylose maize starches on colonic fermentation and apoptotic response to DNA-damage in the colon of rats. Nutr Metab (Lond) 6, 11. 10.1186/1743-7075-6-11.

7. Ramakrishna, B.S., Venkataraman, S., Srinivasan, P., Dash, P., Young, G.P., and Binder, H.J. (2000). Amylase-resistant starch plus oral rehydration solution for cholera. N Engl J Med 342, 308–313. 10.1056/NEJM200002033420502.

8. Snelson, M., Kellow, N.J., and Coughlan, M.T. (2019). Modulation of the Gut Microbiota by Resistant Starch as a Treatment of Chronic Kidney Diseases: Evidence of Efficacy and Mechanistic Insights. Adv Nutr 10, 303–320. 10.1093/advances/nmy068.

9. Maziarz, M.P., Preisendanz, S., Juma, S., Imrhan, V., Prasad, C., and Vijayagopal, P. (2017). Resistant starch lowers postprandial glucose and leptin in overweight adults consuming a moderate-to-high-fat diet: a randomized-controlled trial. Nutr J 16, 14. 10.1186/s12937-017-0235-8.

10. Cerqueira, F.M., Photenhauer, A.L., Pollet, R.M., Brown, H.A., and Koropatkin, N.M. (2020). Starch Digestion by Gut Bacteria: Crowdsourcing for Carbs. Trends Microbiol 28, 95–108. 10.1016/j.tim.2019.09.004.

11. Leitch, E.C., Walker, A.W., Duncan, S.H., Holtrop, G., and Flint, H.J. (2007). Selective colonization of insoluble substrates by human faecal bacteria. Environ Microbiol 9, 667–679. 10.1111/j.1462-2920.2006.01186.x.

12. Ze, X., Duncan, S.H., Louis, P., and Flint, H.J. (2012). Ruminococcus bromii is a keystone species for the degradation of resistant starch in the human colon. ISME J 6, 1535–1543. 10.1038/ismej.2012.4.

13. Rangarajan, A.A., Chia, H.E., Azaldegui, C.A., Olszewski, M.H., Pereira, G.V., Koropatkin, N.M., and Biteen, J.S. (2022). Ruminococcus bromii enables the growth of proximal Bacteroides thetaiotaomicron by releasing glucose during starch degradation. Microbiology (Reading) 168. 10.1099/mic.0.001180.

14. Crost, E.H., Le Gall, G., Laverde-Gomez, J.A., Mukhopadhya, I., Flint, H.J., and Juge, N. (2018). Mechanistic Insights Into the Cross-Feeding of Ruminococcus gnavus and Ruminococcus bromii on Host and Dietary Carbohydrates. Front Microbiol 9, 2558. 10.3389/fmicb.2018.02558.

15. Gopalakrishnan, V., Spencer, C.N., Nezi, L., Reuben, A., Andrews, M.C., Karpinets, T.V., Prieto, P.A., Vicente, D., Hoffman, K., Wei, S.C., et al. (2018). Gut microbiome modulates response to anti-PD-1 immunotherapy in melanoma patients. Science 359, 97–103. 10.1126/science.aan4236.

16. Wang, H., Ainiwaer, A., Song, Y., Qin, L., Peng, A., Bao, H., and Qin, H. (2023). Perturbed gut microbiome and fecal and serum metabolomes are associated with chronic kidney disease severity. Microbiome 11, 3. 10.1186/s40168-022-01443-4.

17. Brushett, S., Gacesa, R., Vich Vila, A., Brandao Gois, M.F., Andreu-Sanchez, S., Swarte, J.C., Klaassen, M.A.Y., Collij, V., Sinha, T., Bolte, L.A., et al. (2023). Gut feelings: the relations between depression, anxiety, psychotropic drugs and the gut microbiome. Gut Microbes 15, 2281360. 10.1080/19490976.2023.2281360.

18. Baxter, N.T., Schmidt, A.W., Venkataraman, A., Kim, K.S., Waldron, C., and Schmidt, T.M. (2019). Dynamics of Human Gut Microbiota and Short-Chain Fatty Acids in Response to Dietary Interventions with Three Fermentable Fibers. mBio 10. 10.1128/mBio.02566-18.

19. Ze, X., Ben David, Y., Laverde-Gomez, J.A., Dassa, B., Sheridan, P.O., Duncan, S.H., Louis, P., Henrissat, B., Juge, N., Koropatkin, N.M., et al. (2015). Unique Organization of Extracellular Amylases into Amylosomes in the Resistant Starch-Utilizing Human Colonic Firmicutes Bacterium Ruminococcus bromii. mBio 6, e01058–15. 10.1128/mBio.01058-15.

20. Tatli, M., Morais, S., Tovar-Herrera, O.E., Bomble, Y.J., Bayer, E.A., Medalia, O., and Mizrahi, I. (2022). Nanoscale resolution of microbial fiber degradation in action. Elife 11. 10.7554/eLife.76523.

21. Artzi, L., Bayer, E.A., and Moraïs, S. (2017). Cellulosomes: bacterial nanomachines for dismantling plant polysaccharides. Nat Rev Microbiol 15, 83–95. 10.1038/nrmicro.2016.164.

22. Mukhopadhya, I., Morais, S., Laverde-Gomez, J., Sheridan, P.O., Walker, A.W., Kelly, W., Klieve, A.V., Ouwerkerk, D., Duncan, S.H., Louis, P., et al. (2018). Sporulation capability and amylosome conservation among diverse human colonic and rumen isolates of the keystone starch-degrader Ruminococcus bromii. Environ Microbiol 20, 324–336. 10.1111/1462-2920.14000.

23. Cockburn, D.W., Kibler, R., Brown, H.A., Duvall, R., Morais, S., Bayer, E., and Koropatkin, N.M. (2021). Structure and substrate recognition by the Ruminococcus bromii amylosome pullulanases. J Struct Biol 213, 107765. 10.1016/j.jsb.2021.107765.

24. Photenhauer, A.L., Villafuerte-Vega, R.C., Cerqueira, F.M., Armbruster, K.M., Marecek, F., Chen, T., Wawrzak, Z., Hopkins, J.B., Vander Kooi, C.W., Janecek, S., et al. (2024). The Ruminococcus bromii amylosome protein Sas6 binds single and double helical alpha-glucan structures in starch. Nat Struct Mol Biol 31, 255–265. 10.1038/s41594-023-01166-6.

25. Cerqueira, F.M., Photenhauer, A.L., Doden, H.L., Brown, A.N., Abdel-Hamid, A.M., Morais, S., Bayer, E.A., Wawrzak, Z., Cann, I., Ridlon, J.M., et al. (2022). Sas20 is a highly flexible starch-binding protein in the Ruminococcus bromii cell-surface amylosome. J Biol Chem 298, 101896. 10.1016/j.jbc.2022.101896.

26. Janecek, S., Marecek, F., MacGregor, E.A., and Svensson, B. (2019). Starch-binding domains as CBM families-history, occurrence, structure, function and evolution. Biotechnol Adv 37, 107451. 10.1016/j.biotechadv.2019.107451.

27. Wangpaiboon, K., Laohawuttichai, P., Kim, S.Y., Mori, T., Nakapong, S., Pichyangkura, R., Pongsawasdi, P., Hakoshima, T., and Krusong, K. (2021). A GH13 alpha-glucosidase from Weissella cibaria uncommonly acts on short-chain maltooligosaccharides. Acta Crystallogr D Struct Biol 77, 1064–1076. 10.1107/S205979832100677X.

28. Cerqueira, F.M. (2022). Biochemical Features of Resistant Starch Degradation by Ruminococcus bromii.

29. El Kaoutari, A., Armougom, F., Gordon, J.I., Raoult, D., and Henrissat, B. (2013). The abundance and variety of carbohydrate-active enzymes in the human gut microbiota. Nat Rev Microbiol 11, 497–504. 10.1038/nrmicro3050.

30. Boraston, A.B., Healey, M., Klassen, J., Ficko-Blean, E., Lammerts van Bueren, A., and Law, V. (2006). A structural and functional analysis of alpha-glucan recognition by family 25 and 26 carbohydrate-binding modules reveals a conserved mode of starch recognition. J Biol Chem 281, 587–598. 10.1074/jbc.M509958200.

31. Miyazaki, K., Martin, J.C., Marinsek-Logar, R., and Flint, H.J. (1997). Degradation and utilization of xylans by the rumen anaerobe Prevotella bryantii (formerly P. ruminicola subsp. brevis) B(1)4. Anaerobe 3, 373–381. 10.1006/anae.1997.0125.

32. Morais, S., Mazor, M., Tovar-Herrera, O., Zehavi, T., Zorea, A., Ifrach, M., Bogumil, D., Brandis, A., Walter, J., Elia, N., et al. (2024). Plasmid-encoded toxin defence mediates mutualistic microbial interactions. Nat Microbiol 9, 108–119. 10.1038/s41564-023-01521-9.

33. Bonn, F., Maass, S., and van Dijl, J.M. (2018). Enrichment of Cell Surface-Associated Proteins in Gram-Positive Bacteria by Biotinylation or Trypsin Shaving for Mass Spectrometry Analysis. Methods Mol Biol 1841, 35–43. 10.1007/978-1-4939-8695-8_4.

34. Otto, A., Maass, S., Bonn, F., Buttner, K., and Becher, D. (2017). An Easy and Fast Protocol for Affinity Bead-Based Protein Enrichment and Storage of Proteome Samples. Methods Enzymol 585, 1–13. 10.1016/bs.mie.2016.09.012.

35. Bonn, F., Bartel, J., Buttner, K., Hecker, M., Otto, A., and Becher, D. (2014). Picking vanished proteins from the void: how to collect and ship/share extremely dilute proteins in a reproducible and highly efficient manner. Anal Chem 86, 7421–7427. 10.1021/ac501189j.

36. Cox, J., and Mann, M. (2008). MaxQuant enables high peptide identification rates, individualized p.p.b.-range mass accuracies and proteome-wide protein quantification. Nat Biotechnol 26, 1367–1372. 10.1038/nbt.1511.

37. Cox, J., Hein, M.Y., Luber, C.A., Paron, I., Nagaraj, N., and Mann, M. (2014). Accurate proteome-wide label-free quantification by delayed normalization and maximal peptide ratio extraction, termed MaxLFQ. Mol Cell Proteomics 13, 2513–2526. 10.1074/mcp.M113.031591.

38. Cox, J., Neuhauser, N., Michalski, A., Scheltema, R.A., Olsen, J.V., and Mann, M. (2011). Andromeda: a peptide search engine integrated into the MaxQuant environment. J Proteome Res 10, 1794–1805. 10.1021/pr101065j.

39. Tyanova, S., and Cox, J. (2018). Perseus: A Bioinformatics Platform for Integrative Analysis of Proteomics Data in Cancer Research. Methods Mol Biol 1711, 133–148. 10.1007/978-1-4939-7493-1_7.

40. Zheng, J., Ge, Q., Yan, Y., Zhang, X., Huang, L., and Yin, Y. (2023). dbCAN3: automated carbohydrate-active enzyme and substrate annotation. Nucleic Acids Res 51, W115–W121. 10.1093/nar/gkad328.

41. Yu, N.Y., Wagner, J.R., Laird, M.R., Melli, G., Rey, S., Lo, R., Dao, P., Sahinalp, S.C., Ester, M., Foster, L.J., et al. (2010). PSORTb 3.0: improved protein subcellular localization prediction with refined localization subcategories and predictive capabilities for all prokaryotes. Bioinformatics 26, 1608–1615. 10.1093/bioinformatics/btq249.

42. Barak, Y., Handelsman, T., Nakar, D., Mechaly, A., Lamed, R., Shoham, Y., and Bayer, E.A. (2005). Matching fusion protein systems for affinity analysis of two interacting families of proteins: the cohesin-dockerin interaction. J Mol Recognit 18, 491–501. 10.1002/jmr.749.

43. Punjani, A., and Fleet, D.J. (2023). 3DFlex: determining structure and motion of flexible proteins from cryo-EM. Nat Methods 20, 860–870. 10.1038/s41592-023-01853-8.

44. Jumper, J., Evans, R., Pritzel, A., Green, T., Figurnov, M., Ronneberger, O., Tunyasuvunakool, K., Bates, R., Zidek, A., Potapenko, A., et al. (2021). Highly accurate protein structure prediction with AlphaFold. Nature 596, 583–589. 10.1038/s41586-021-03819-2.

45. Meng, E.C., Goddard, T.D., Pettersen, E.F., Couch, G.S., Pearson, Z.J., Morris, J.H., and Ferrin, T.E. (2023). UCSF ChimeraX: Tools for structure building and analysis. Protein Sci 32, e4792. 10.1002/pro.4792.

46. Terwilliger, T.C., Liebschner, D., Croll, T.I., Williams, C.J., McCoy, A.J., Poon, B.K., Afonine, P.V., Oeffner, R.D., Richardson, J.S., Read, R.J., et al. (2024). AlphaFold predictions are valuable hypotheses and accelerate but do not replace experimental structure determination. Nat Methods 21, 110–116. 10.1038/s41592-023-02087-4.

47. Croll, T.I. (2018). ISOLDE: a physically realistic environment for model building into low-resolution electron-density maps. Acta Crystallogr D Struct Biol 74, 519–530. 10.1107/S2059798318002425.

48. Cockburn, D., and Koropatkin, N. (2015). Product Analysis of Starch Active Enzymes by TLC. Bio-protocol 5, e1621. 10.21769/BioProtoc.1621.

49. Arnal, G., Attia, M.A., Asohan, J., Lei, Z., Golisch, B., and Brumer, H. (2023). A Low-Volume, Parallel Copper-Bicinchoninic Acid (BCA) Assay for Glycoside Hydrolases. Methods Mol Biol 2657, 3–14. 10.1007/978-1-0716-3151-5_1.

50. Wolff, G., Limpens, R., Zheng, S., Snijder, E.J., Agard, D.A., Koster, A.J., and Barcena, M. (2019). Mind the gap: Micro-expansion joints drastically decrease the bending of FIB-milled cryo-lamellae. J Struct Biol 208, 107389. 10.1016/j.jsb.2019.09.006.

51. Mastronarde, D.N. (2003). SerialEM: A Program for Automated Tilt Series Acquisition on Tecnai Microscopes Using Prediction of Specimen Position. Microsc Microanal 9, 1182–1183. 10.1017/S1431927603445911.

52. Eisenstein, F., Yanagisawa, H., Kashihara, H., Kikkawa, M., Tsukita, S., and Danev, R. (2023). Parallel cryo electron tomography on in situ lamellae. Nat Methods 20, 131–138. 10.1038/s41592-022-01690-1.

53. Zheng, S.Q., Palovcak, E., Armache, J.P., Verba, K.A., Cheng, Y., and Agard, D.A. (2017). MotionCor2: anisotropic correction of beam-induced motion for improved cryo-electron microscopy. Nat Methods 14, 331–332. 10.1038/nmeth.4193.

54. Kremer, J.R., Mastronarde, D.N., and McIntosh, J.R. (1996). Computer visualization of three-dimensional image data using IMOD. J Struct Biol 116, 71–76. 10.1006/jsbi.1996.0013.

55. Zheng, S., Wolff, G., Greenan, G., Chen, Z., Faas, F.G.A., Barcena, M., Koster, A.J., Cheng, Y., and Agard, D.A. (2022). AreTomo: An integrated software package for automated marker-free, motion-corrected cryo-electron tomographic alignment and reconstruction. J Struct Biol X 6, 100068. 10.1016/j.yjsbx.2022.100068.

56. Buchholz, T.O., Krull, A., Shahidi, R., Pigino, G., Jekely, G., and Jug, F. (2019). Content-aware image restoration for electron microscopy. Methods Cell Biol 152, 277–289. 10.1016/bs.mcb.2019.05.001.

57. Wiedemann, S., and Heckel, R. (2024). A deep learning method for simultaneous denoising and missing wedge reconstruction in cryogenic electron tomography. Nat Commun 15, 8255. 10.1038/s41467-024-51438-y.

58. Lamm, L., Zufferey, S., Righetto, R.D., Wietrzynski, W., Yamauchi, K.A., Burt, A., Liu, Y., Zhang, H., Martinez-Sanchez, A., Ziegler, S., et al. (2024). MemBrain v2: an end-to-end tool for the analysis of membranes in cryo-electron tomography. bioRxiv, 2024.01.05.574336. 10.1101/2024.01.05.574336.

59. Jin, W., Zhou, Y., and Bartesaghi, A. (2024). Accurate size-based protein localization from cryo-ET tomograms. J Struct Biol X 10, 100104. 10.1016/j.yjsbx.2024.100104.

60. Chaillet, M.L., van der Schot, G., Gubins, I., Roet, S., Veltkamp, R.C., and Forster, F. (2023). Extensive Angular Sampling Enables the Sensitive Localization of Macromolecules in Electron Tomograms. Int J Mol Sci 24. 10.3390/ijms241713375.

61. alisterburt, Dimitry Tegunov, tegunovd, and Moritz Wachsmuth-Melm (2025). warpem/warp: v2.0.0dev32. Version v2.0.0dev32 (Zenodo). 10.5281/ZENODO.13982246 10.5281/ZENODO.13982246.

62. Burt, A., Toader, B., Warshamanage, R., von Kügelgen, A., Pyle, E., Zivanov, J., Kimanius, D., Bharat, T.A.M., and Scheres, S.H.W. (2024). An image processing pipeline for electron cryo-tomography in RELION-5. FEBS Open Bio 14, 1788–1804. 10.1002/2211-5463.13873.

63. Tegunov, D., Xue, L., Dienemann, C., Cramer, P., and Mahamid, J. (2021). Multi-particle cryo-EM refinement with M visualizes ribosome-antibiotic complex at 3.5 A in cells. Nat Methods 18, 186–193. 10.1038/s41592-020-01054-7.

64. Ermel, U.H., Arghittu, S.M., and Frangakis, A.S. (2022). ArtiaX: An electron tomography toolbox for the interactive handling of sub-tomograms in UCSF ChimeraX. Protein Sci 31, e4472. 10.1002/pro.4472.

65. Perez-Riverol, Y., Bai, J., Bandla, C., Garcia-Seisdedos, D., Hewapathirana, S., Kamatchinathan, S., Kundu, D.J., Prakash, A., Frericks-Zipper, A., Eisenacher, M., et al. (2022). The PRIDE database resources in 2022: a hub for mass spectrometry-based proteomics evidences. Nucleic Acids Res 50, D543–D552. 10.1093/nar/gkab1038.

