## Supplementary material for "Spatial constraints drive amylosome-mediated resistant starch degradation by *Ruminococcus bromii* in the human colon": Document S1, with Figs. S1 - S15

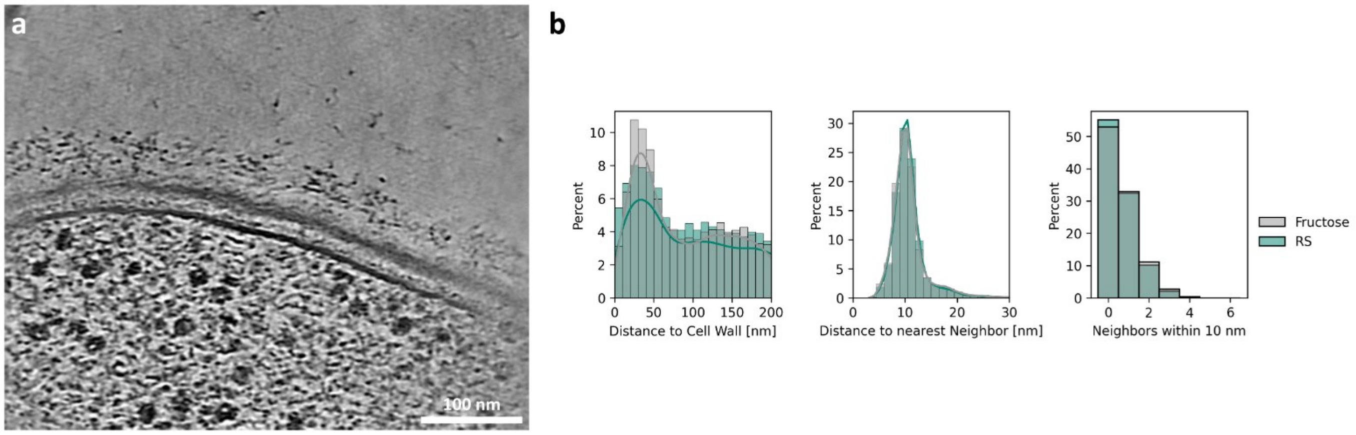


**Figure S1: Representative tomogram of *R. bromii* grown on fructose**

**(a)** High-magnification tomogram of the edge of *R. bromii* grown on fructose shows the amylosome around the cell. The architecture of the amylosomes is visible as globular densities anchored to the cell wall through elongated linkers. **(b)** Plots showing the protein density in the amylosome outside the cell, compared between cells grown on RS and on fructose. No significant changes in the distribution are observed.

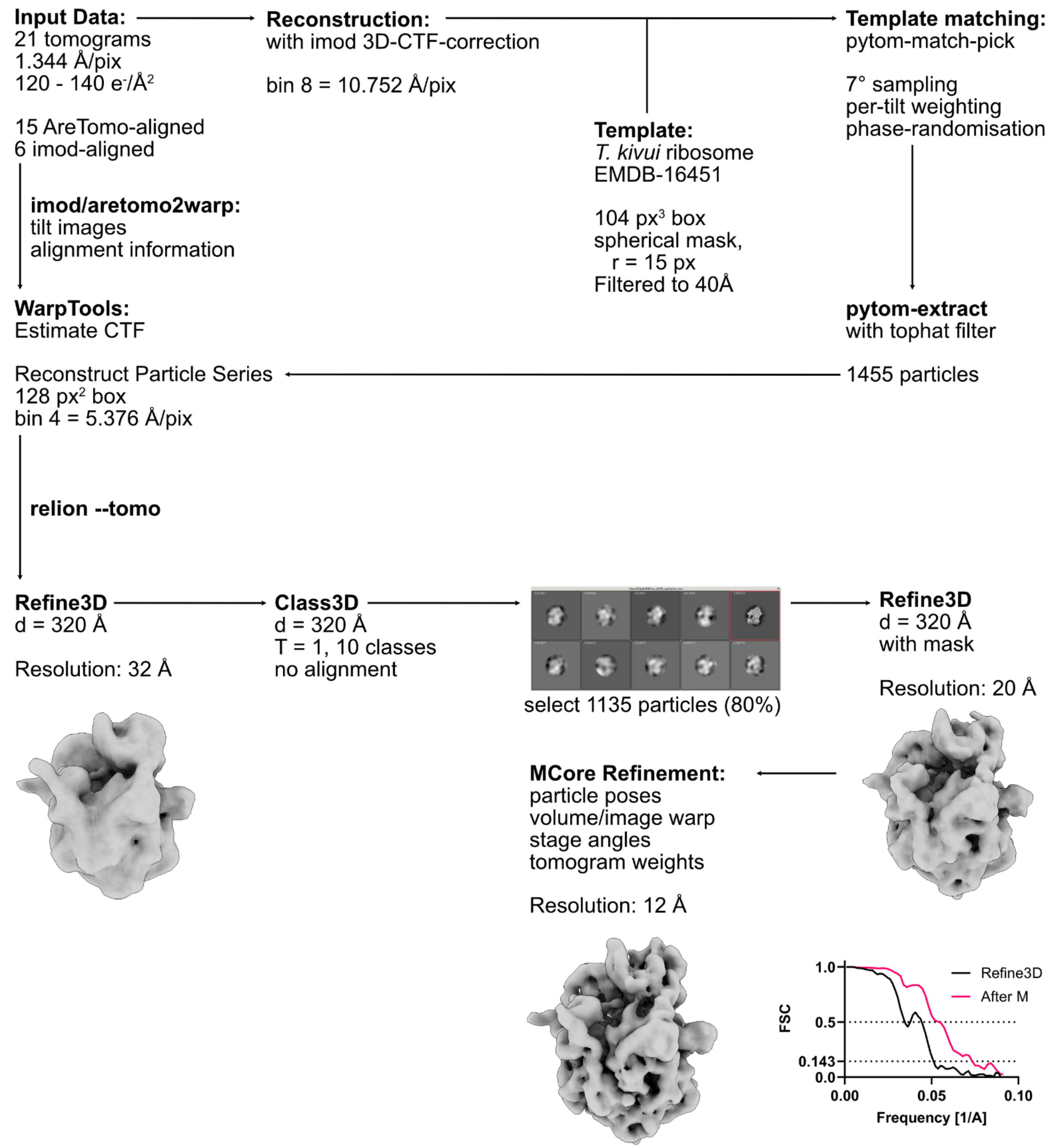


**Figure S2: Subtomogram averaging of ribosomes**

The data processing scheme for subtomogram averaging of the *R. bromii* ribosome. A subset of 21 tomograms were selected for good alignment quality and visibility of membrane cross-sections. They were reconstructed at bin 8 with 3D-CTF-correction for template matching against EMDB-16451 to retrieve initial ribosome coordinates. Tilt images and tilt alignment information were exported from AreTomo or imod to WarpTools using aretomo2warp and imod2warp, respectively, for CTF estimation and particle series reconstruction. Processing in Relion 5 Tomography mode yielded a map containing 80% of initial picks, refining to a resolution of 20 Å. Resolution was further improved by refinement using the software M, as can be seen both from the resulting density map and the Fourier shell correlation (FSC) curve for Relion and M refinement compared.

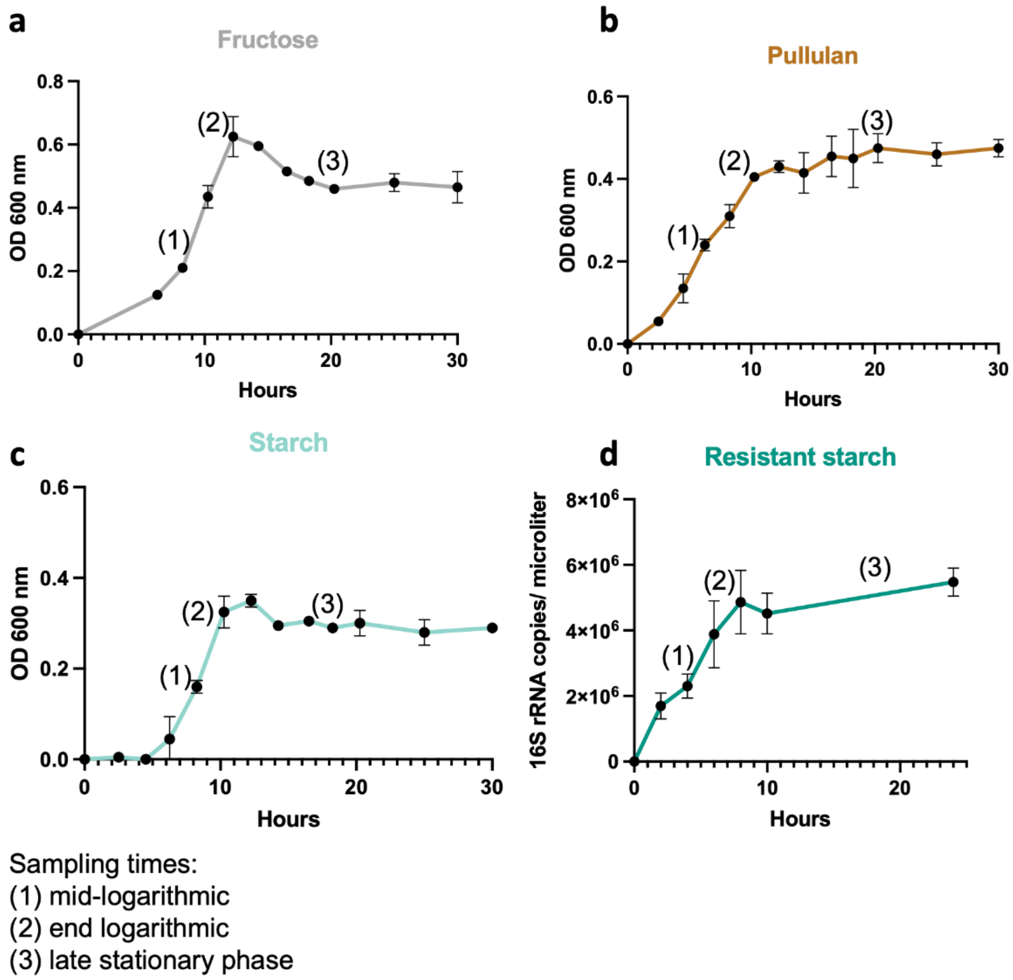


**Figure S3: Growth curves on the selected four carbon sources.**

Growth (monitored by OD 600nm) is shown for *R. bromii* in M2 medium, containing 0.2% of the carbohydrate indicated: **(a)** fructose, **(b)** pullulan, **(c)** starch and **(d)** resistant starch. For resistant starch, growth was monitored by 16S qPCR instead. Sampling times are indicated by numbers on each curve.

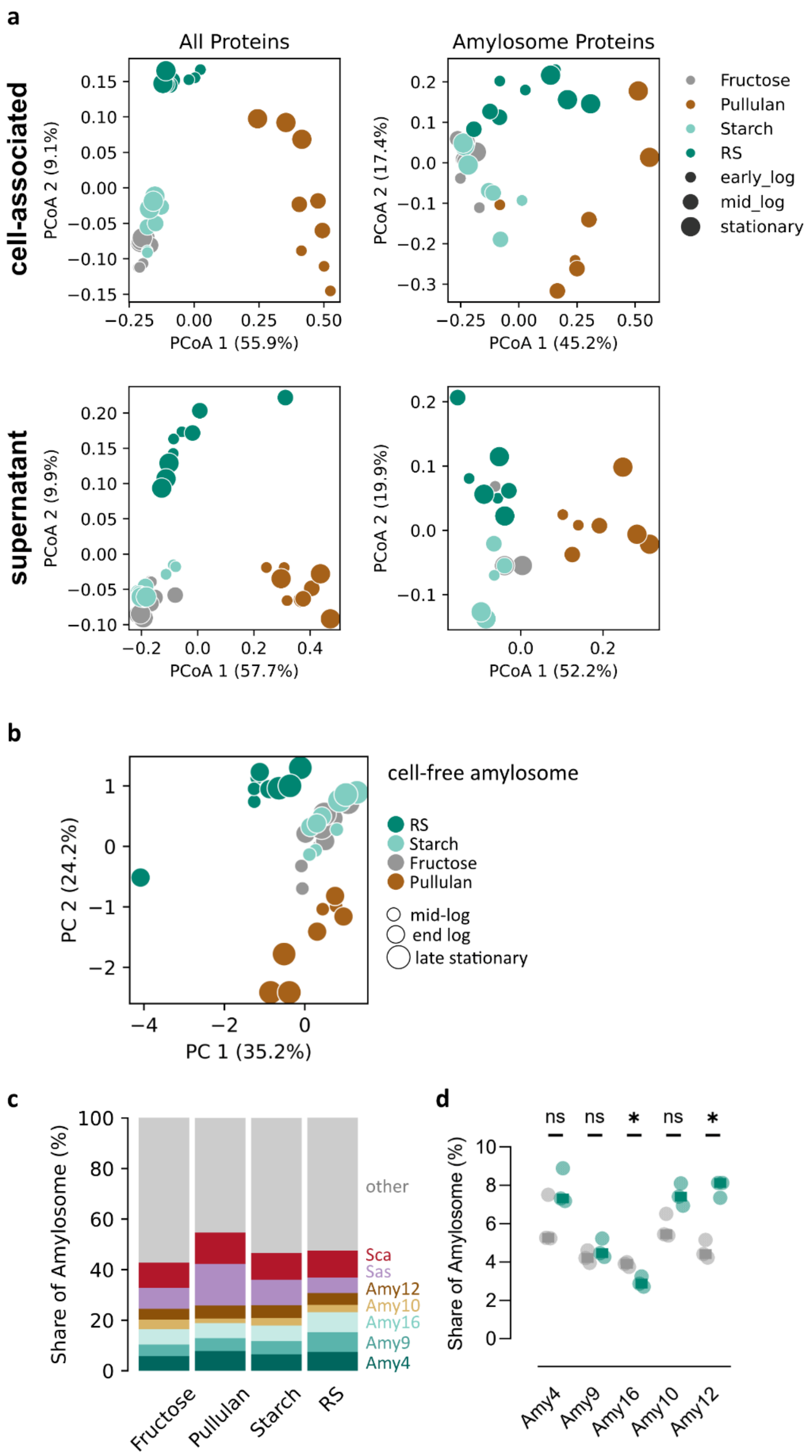


**Figure S4: Functional adaptation of the proteome as a function of the carbon source.**

**(a)** Principal coordinates analysis (PCoA) on the presence and absence of proteins in cell-associated (top) and supernatant (bottom) data, separated into full proteome (left) and amylosome-associated proteins (right). **(b)** A PCA of the protein abundances in the cell-free amylosome shows weak clustering by carbon source (ANOSIM R = 0.52, p = 0.0001). **(c)** Analysis of the average shares of each amylosome component by carbon source. (d) Among the CAZymes, only the shares of Amy16 and Amy12 differ between fructose- and RS-grown cultures at the late stationary phase timepoint (unpaired t-test with Holm-Sidak multiple hypothesis testing correction).

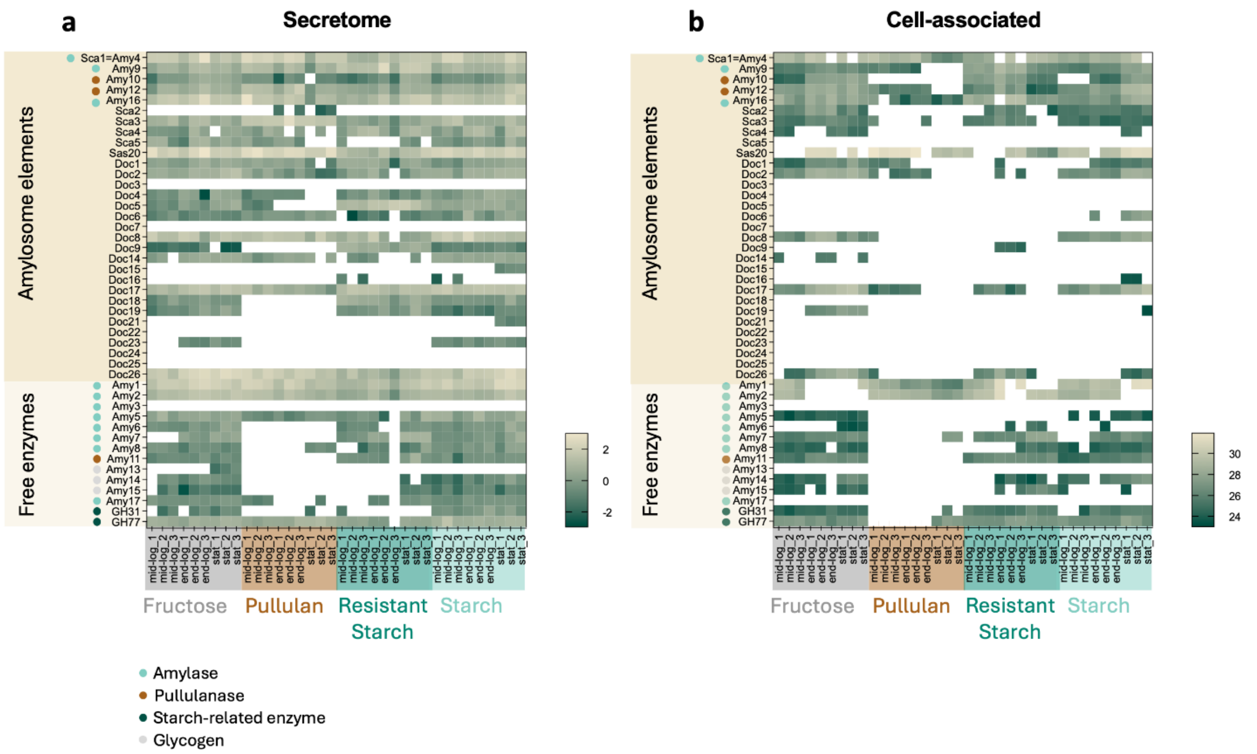


**Figure S5: Proteomic analysis of the amylosome and starch-related enzymatic proteins.**

Heatmap of the secretome **(a)** and cell-associated proteome **(b)** for the various amylosome components is given for the four carbon sources and three sampling times. The scale indicates either Z-Score normalized log 2 transformed LFQ intensity (a) or log 2 transformed LFQ intensity (b). White colors designate that the proteins were not identified under this condition.

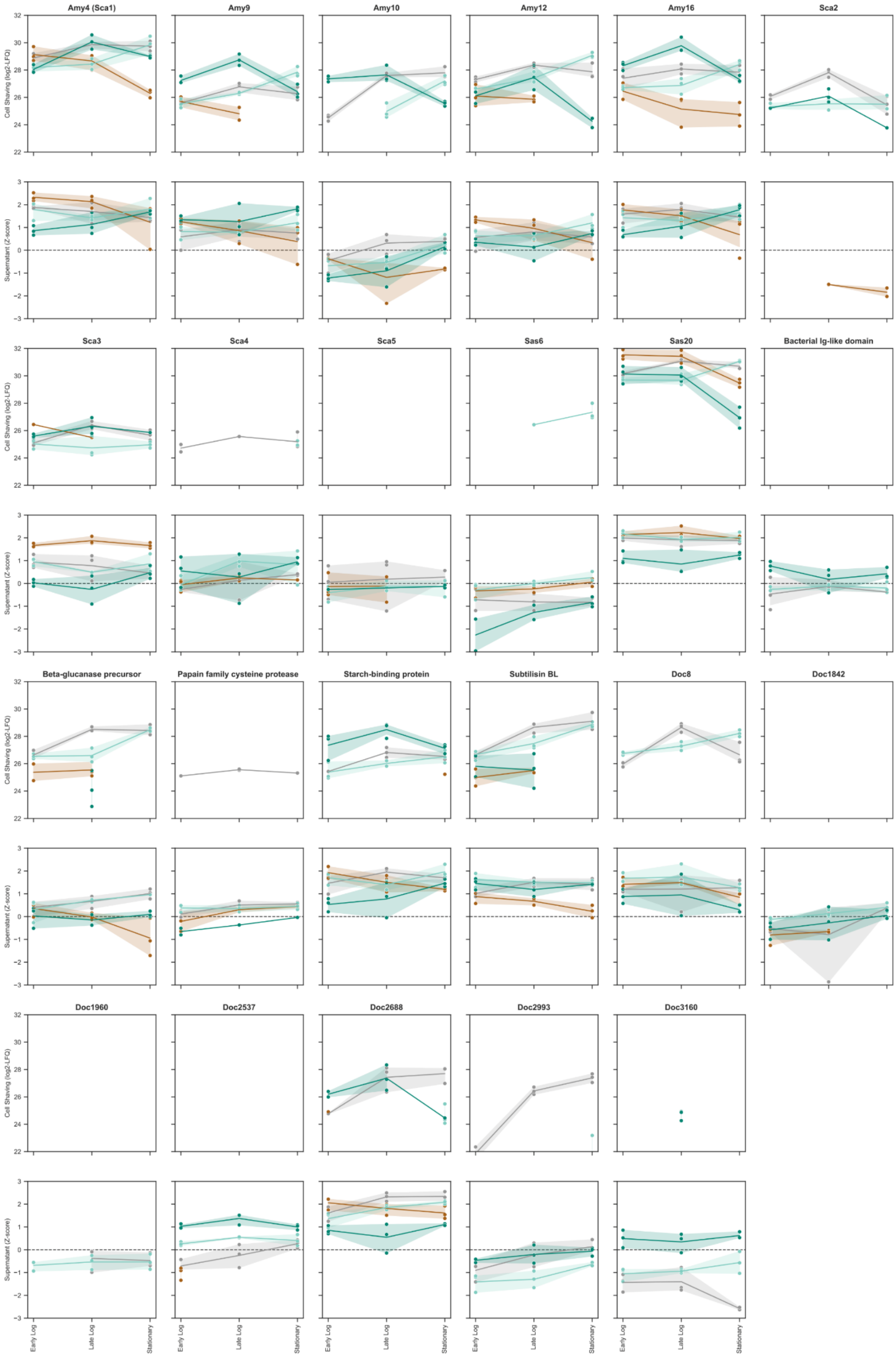


**Figure S6: Expression of the major amylosome components over time.**

Shown are the quantitative values for all amylosome-associated proteins, which were identified in at least two timepoints in either cell-associated or supernatant proteome. First row: Log2-transformed LFQ values. Second row: Z-score normalized abundance for supernatant proteome. Mean and 95% CI are shown.

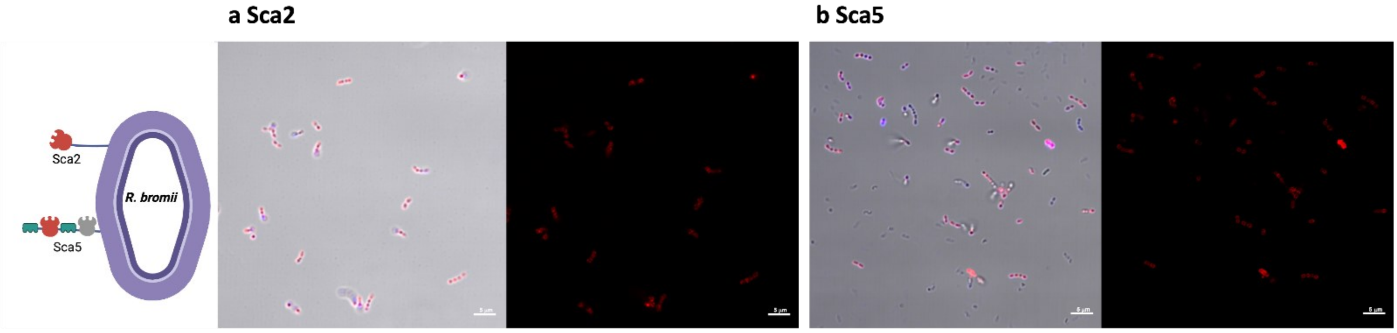


**Figure S7: Detection of cell-wall-anchored scaffoldins.**

Immunofluorescence confocal microscopy images of *R. bromii* confirm the presence of scaffoldins anchored to the cell-wall, using specific primary antibodies directed towards Sca2 **(a)** and Sca5 **(b)**.

Schematic created using biorender.com and available at https://biorender.com/q38pj4a

a

b

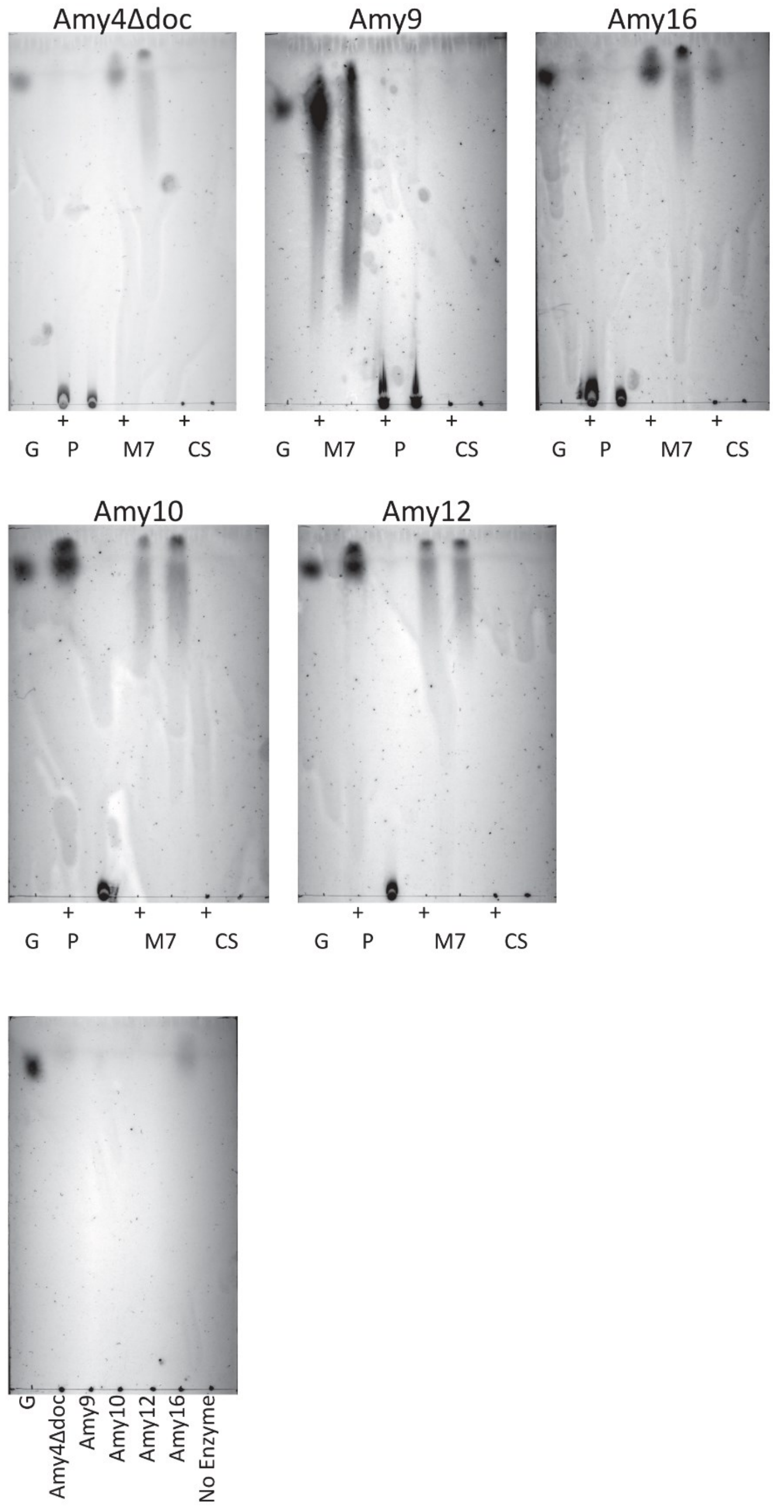


**Figure S8: CAZyme substrate preferences were determined by thin-layer chromatography (TLC).**

**(a)** A standard panel of three carbon sources, pullulan (P), maltoheptaose (M7) and cornstarch (CS) were incubated for 1 h at 37°C either in the presence (+) or absence of each purified enzyme. Substrate transformation was assessed using TLC, run against a 10 mM glucose standard (G). **(b)** Additionally, RS HiMaize 958 was incubated for 3 h at 37°C while shaking with each enzyme or in absence of enzyme. For each reaction, a representative TLC plate is shown.

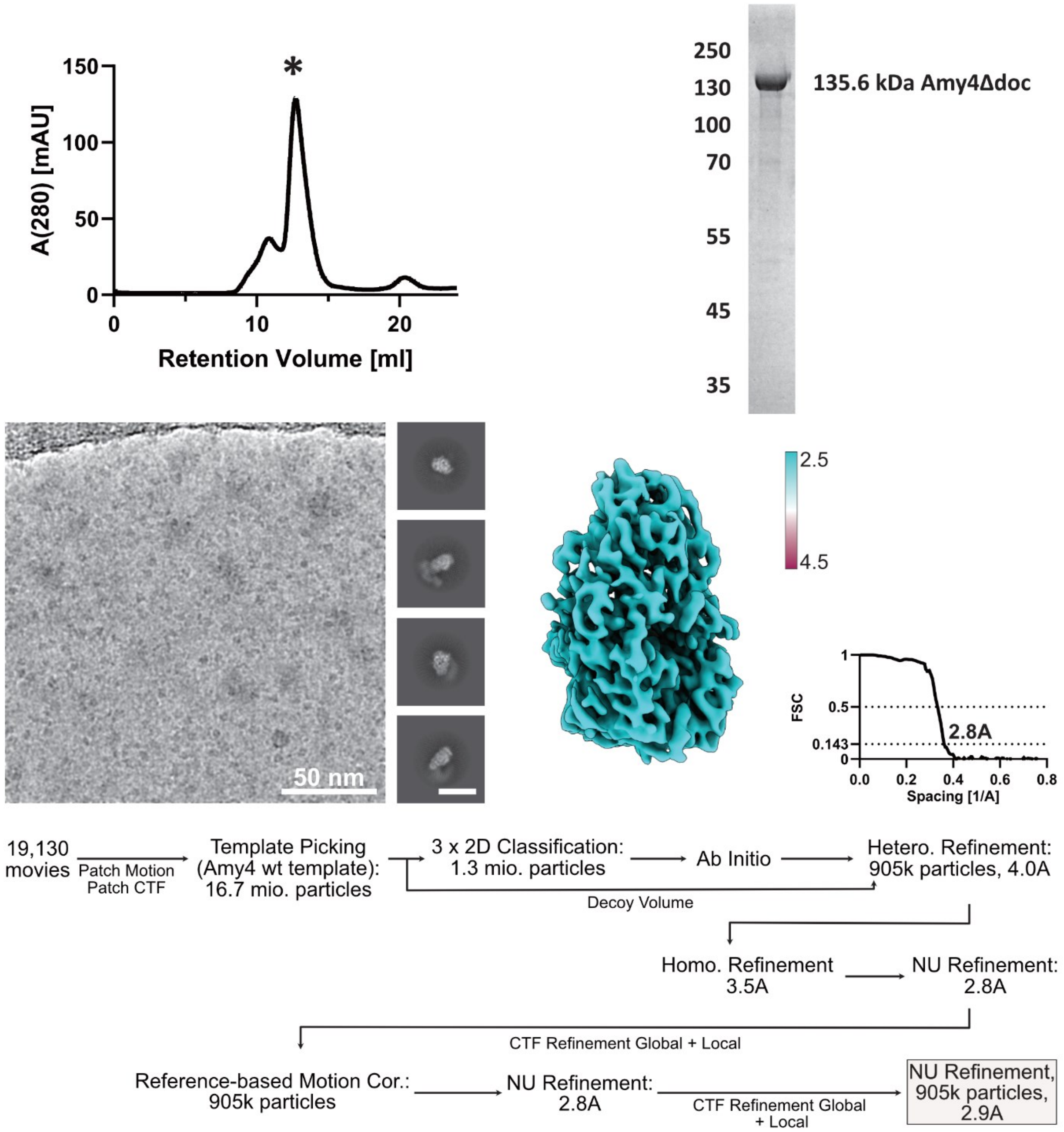


**Figure S9: Amy4Δdoc purification and structure determination.**

Representative SEC chromatogram, selected peak is indicated by an asterisk. Representative SDS-PAGE gel showing purity of the protein, with indicated molecular weight markers (kDa). Micrograph showing particle distribution (scale bar 50 nm), and representative 2D class averages (scale bar 10 nm). Final map colored by local resolution and final masked FSC curve with the resolution at 0.143 indicated. Below, the processing workflow from data to final structure.

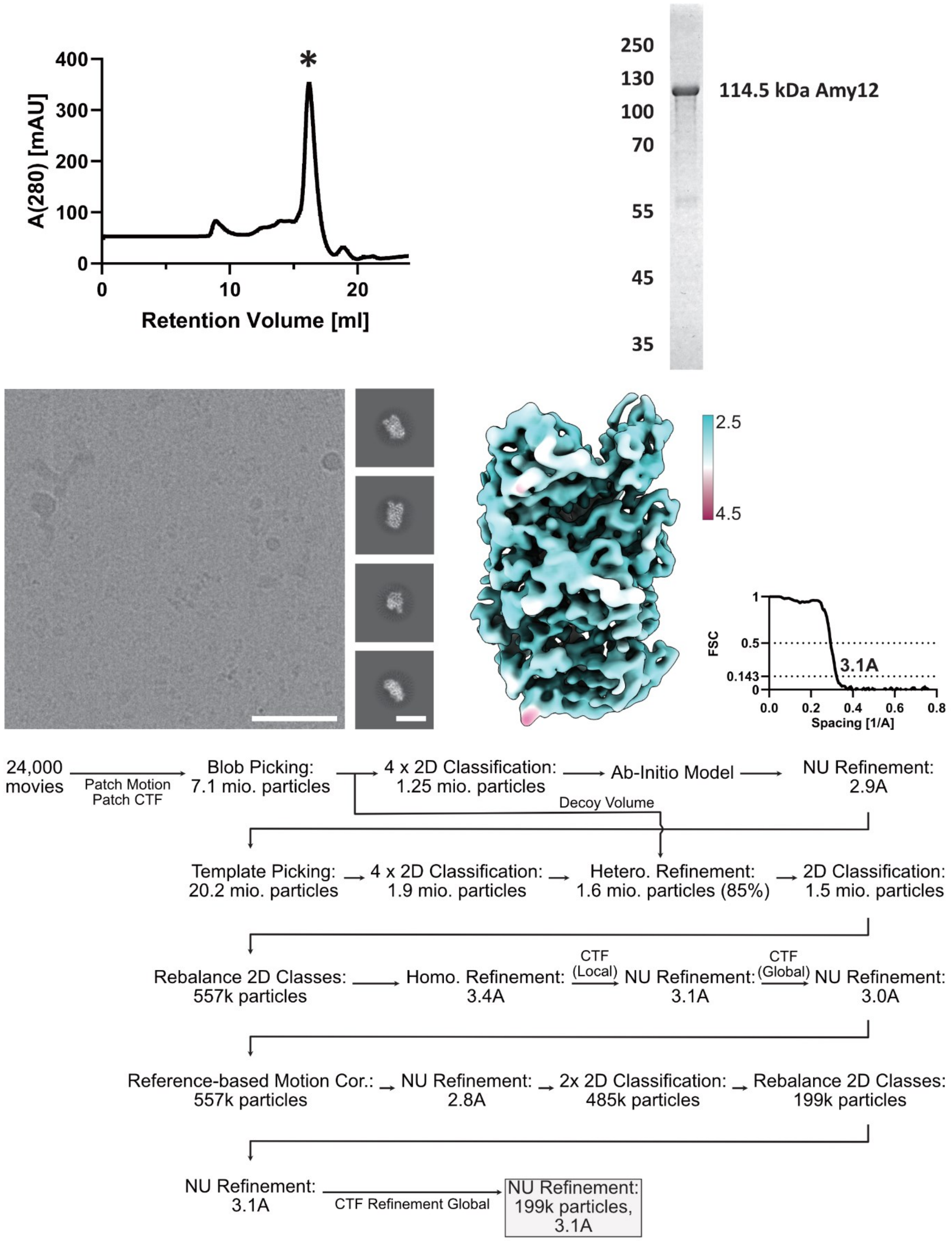


**Figure S10: Amy10 purification and structure determination.**

Representative SEC chromatogram, selected peak indicated by an asterisk. Representative SDS-PAGE gel showing purity of the protein, with indicated molecular weight markers. Micrograph showing particle distribution (scale bar 50 nm), and representative 2D class averages (scale bar 10 nm). Final map colored by local resolution and final masked FSC curve with the resolution at 0.143 indicated. Below, the processing workflow from data to final structure.

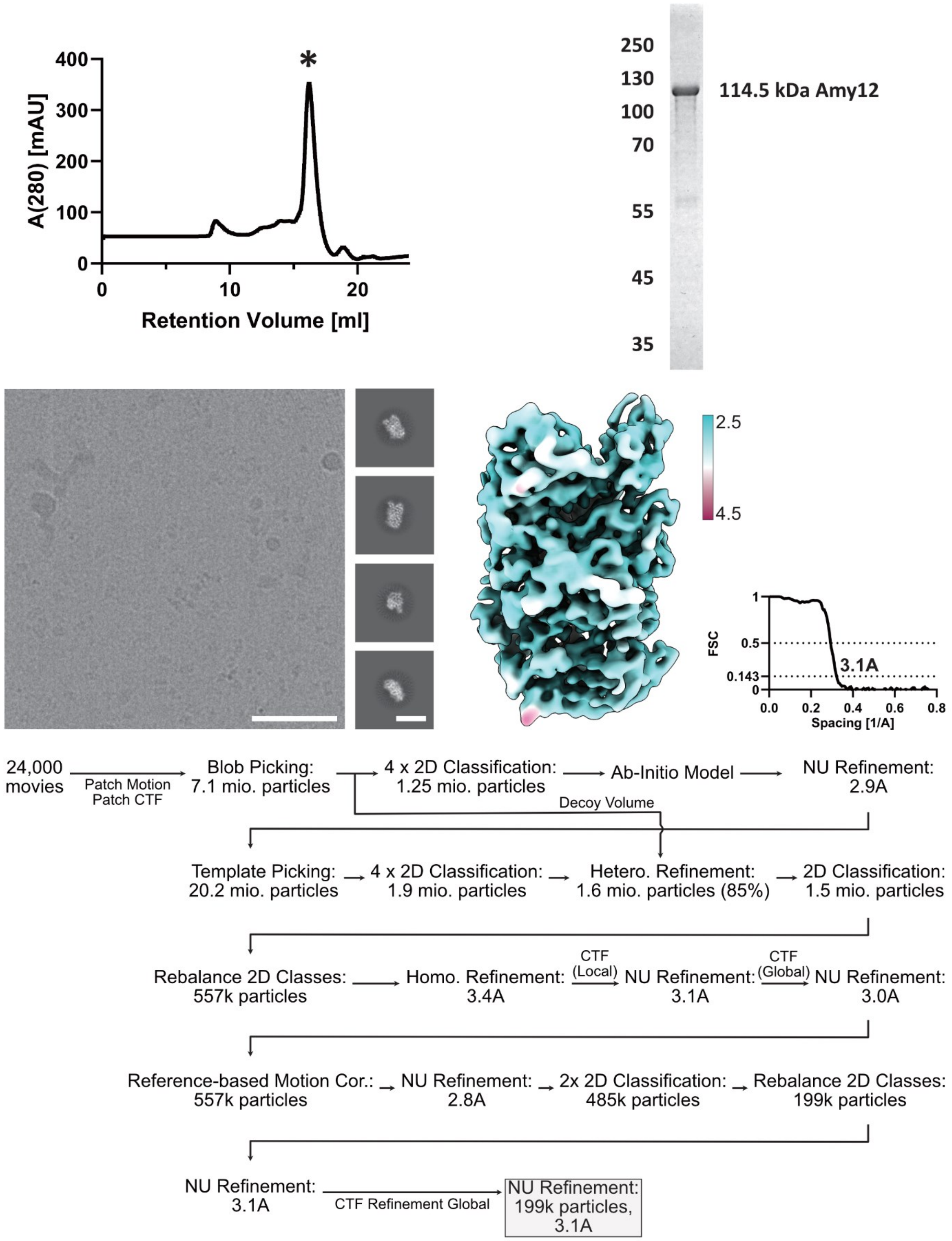


**Figure S11: Amy12 purification and structure determination.**

Representative SEC chromatogram, selected peak indicated by an asterisk. Representative SDS-PAGE gel showing purity of the protein, with indicated molecular weight markers. Micrograph showing particle distribution (scale bar 50 nm), and representative 2D class averages (scale bar 10 nm). Final map colored by local resolution and final masked FSC curve with the resolution at 0.143 indicated. Below, the processing workflow from data to final structure.

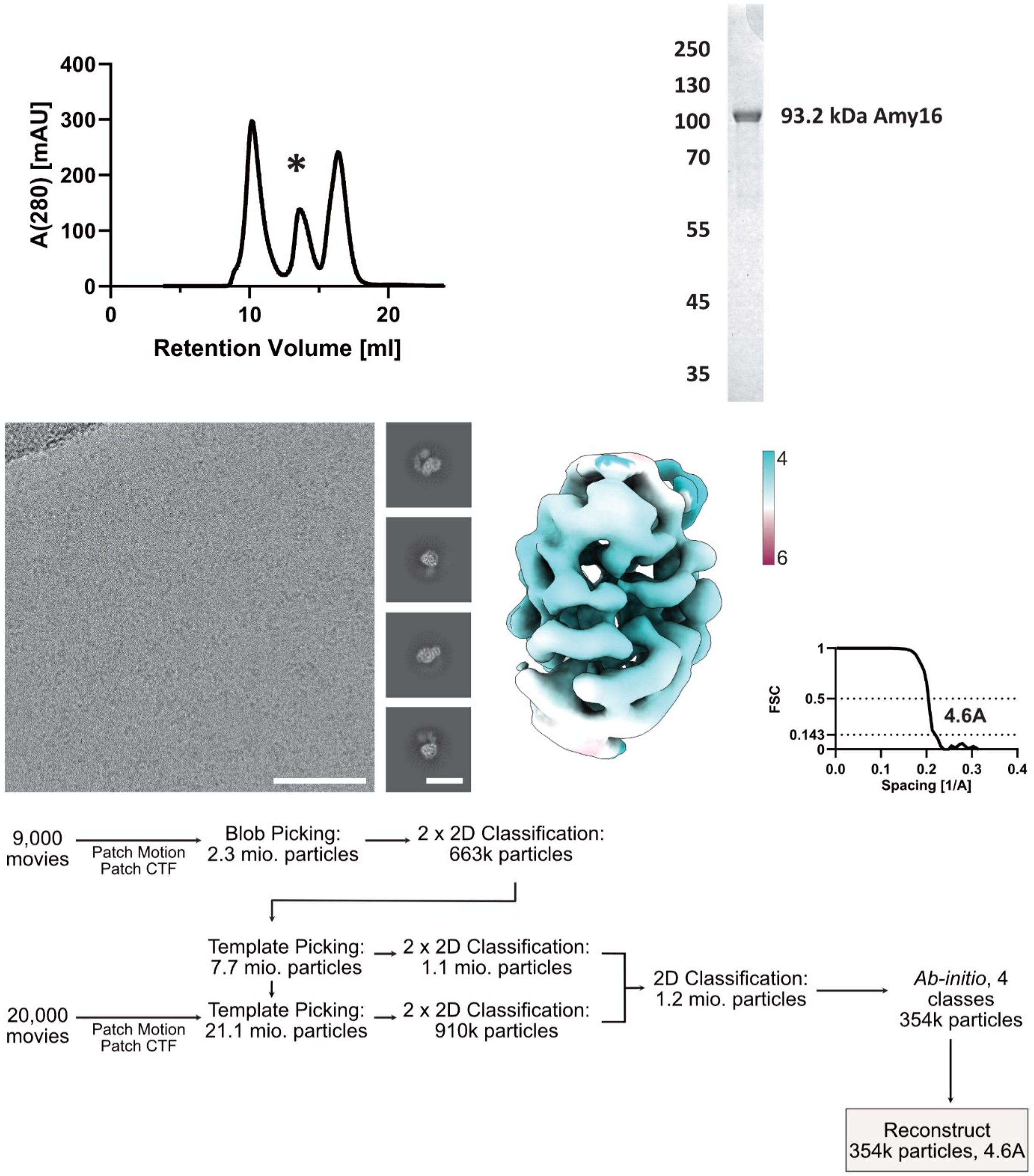


**Figure S12: Amy16 purification and structure determination.**

Representative SEC chromatogram, selected peak indicated by an asterisk. Representative SDS-PAGE gel showing purity of the protein, with indicated molecular weight markers. Micrograph showing particle distribution (scale bar 50 nm), and representative 2D class averages (scale bar 10 nm). Final map colored by local resolution and final masked FSC curve with the resolution at 0.143 indicated. Below, the processing workflow from data to final structure.

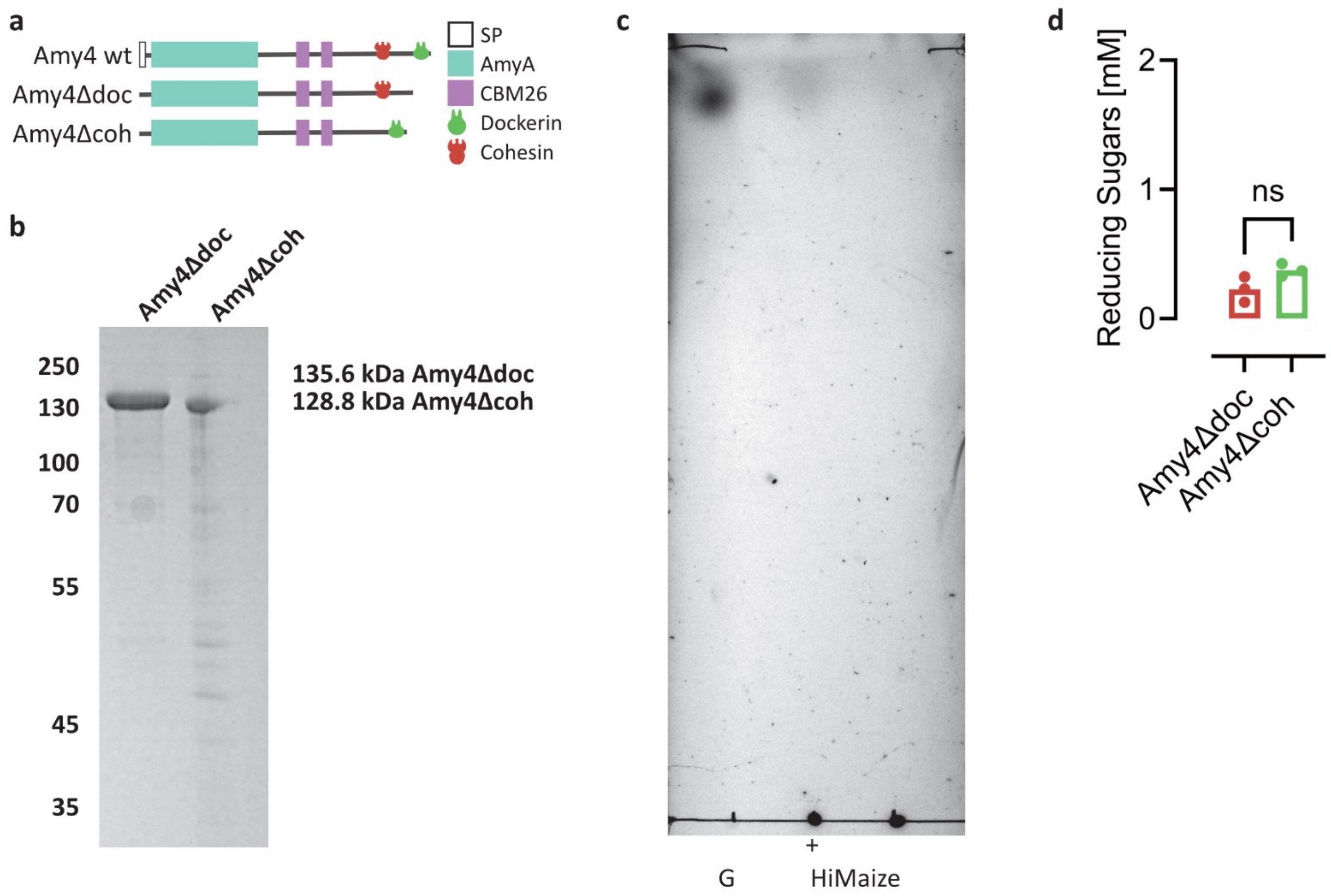


**Figure S13: Purity and activity of Amy4Δdoc and Amy4Δcoh.**

**(a)** A schematic showing the domain organization of wild type Amy4 and the two truncation mutants Amy4Δdoc and Amy4Δcoh, retain either the dockerin or the cohesin domain. **(b)** An SDS-PAGE gel showing the purity of both proteins after SEC. **(c)** A TLC of Amy4Δcoh incubated with HiMaize 958 RS for 3 h. **(d)** A BCA-based quantification of the reducing sugars released by Amy4Δdoc and Amy4Δcoh from HiMaize RS after 3 h (3 technical replicates, unpaired t-test).

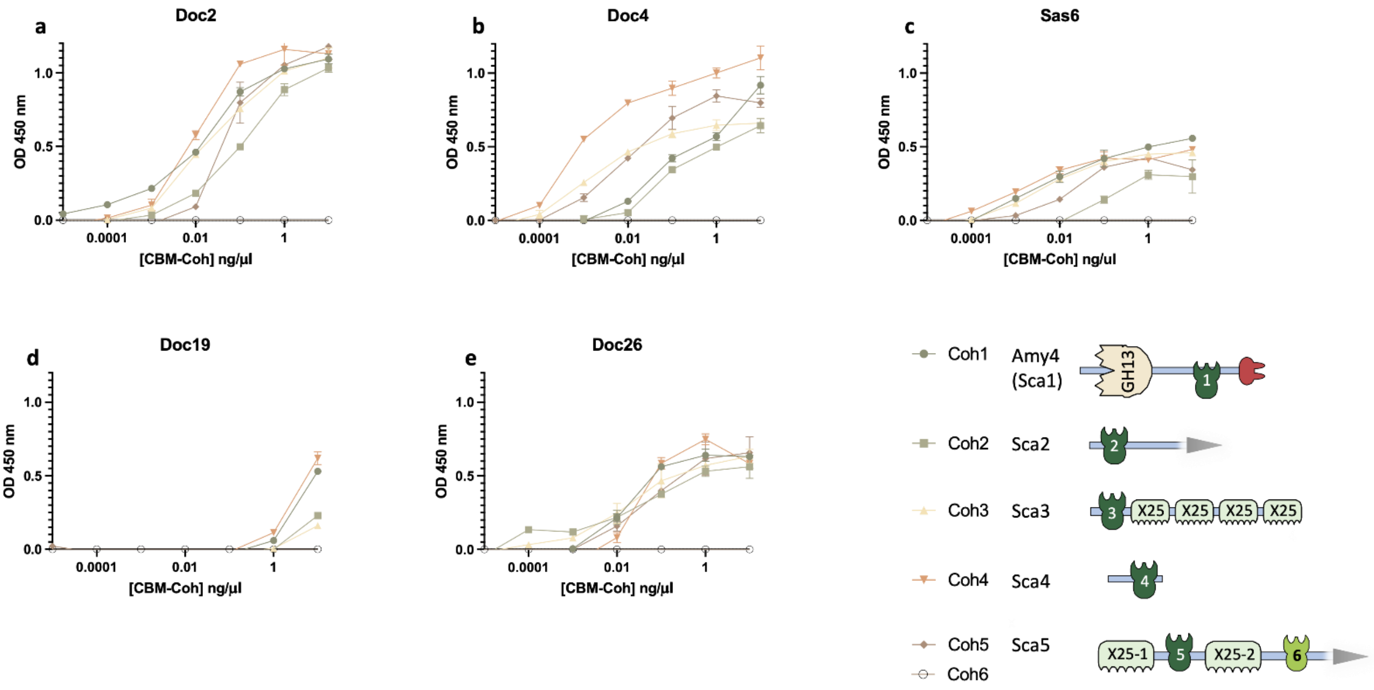


**Figure S14: Cohesin interactions of recombinant dockerins from *R. bromii*.**

Dockerin-containing enzymes detected in proteomic analyses and for which interactions were not previously reported (Supplementary Table 1) were tested for their interactions as represented by absorbance at OD 450 nm with the six different cohesins of *R. bromii* in ELISA experiments. Error bars indicate the standard deviations from the means of the results determined for triplicate samples from one experiment. Each experiment was replicated at least twice.

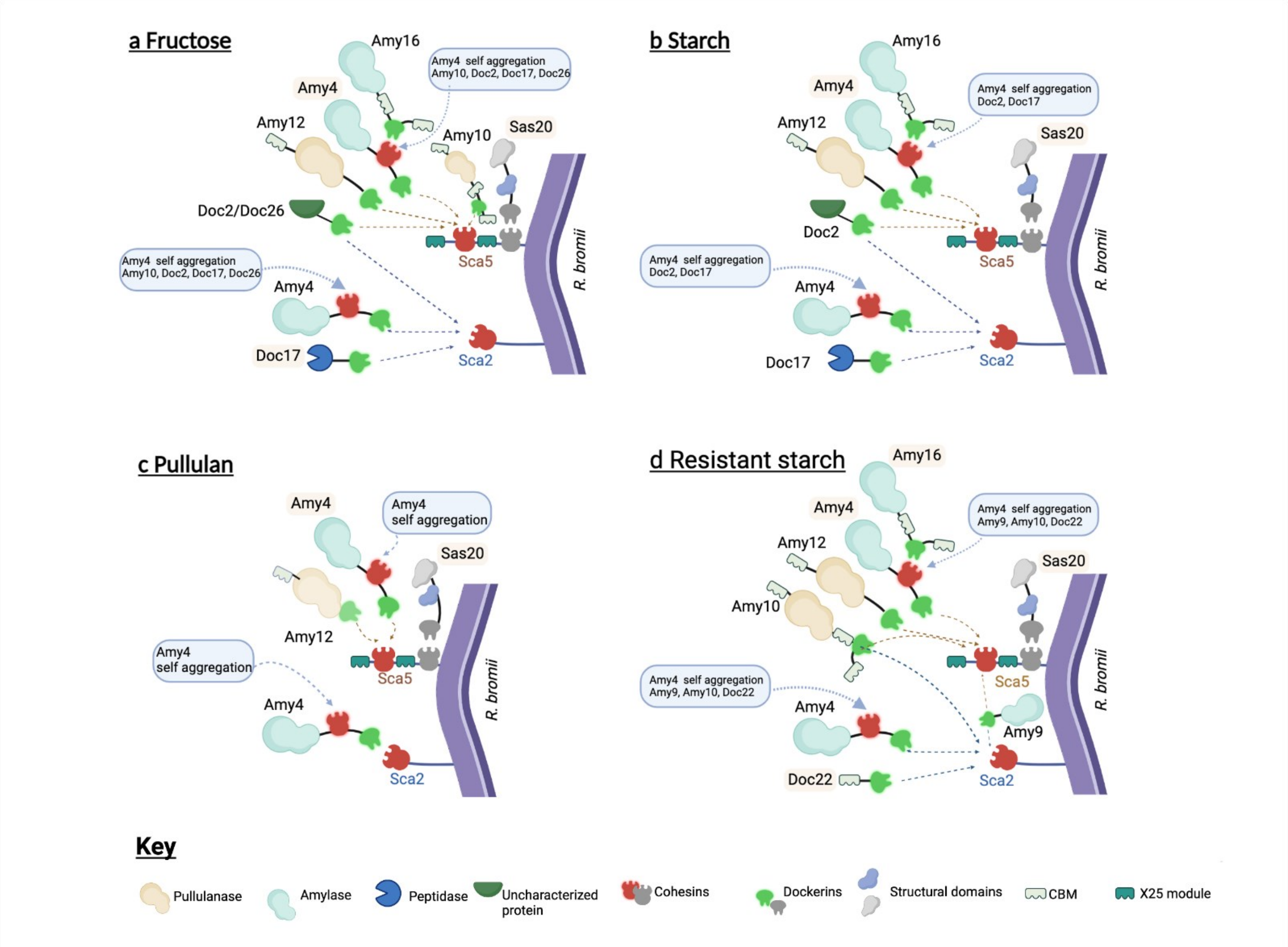


**Figure S15: Updated model for cell-associated amylosomes**

Schematic shows the different amylosome compositions expressed on fructose **(a)**, starch **(b)**, pullulan **(c)** and resistant starch **(d)** in *R. bromii* L2–63. The schematic architecture of each expressed component, as well as binding specificities, is shown in each panel. Only proteins constituting at least 4% of the amylosome content are represented (1% for pullulan), and protein names constituting at least 10% of the amylosome content are highlighted. The schemes depict data from the late stationary phase, except for pullulan, which is shown at the end of the logarithmic phase, where Amy12 was still detected. Sca2 and Sca5 scaffoldins also contain sortase motifs involved in covalent binding to the cell wall, which are not represented.

Panels created with BioRender and available under the following URLs: https://biorender.com/6wcpbyv (fructose), https://biorender.com/eta13cf (starch), https://biorender.com/yjeojtk (pullulan), https://biorender.com/2tseqvv (resistant starch)
